# Inferring the ancestry of everyone

**DOI:** 10.1101/458067

**Authors:** Jerome Kelleher, Yan Wong, Patrick K. Albers, Anthony W. Wohns, Gil McVean

## Abstract

A central problem in evolutionary biology is to infer the full genealogical history of a set of DNA sequences. This history contains rich information about the forces that have influenced a sexually reproducing species. However, existing methods are limited: the most accurate is unable to cope with more than a few dozen samples. With modern genetic data sets rapidly approaching millions of genomes, there is an urgent need for efficient inference methods to exploit such rich resources. We introduce an algorithm to infer whole-genome history which has comparable accuracy to the state-of-the-art but can process around four orders of magnitude more sequences. Additionally, our method results in an “evolutionary encoding” of the original sequence data, enabling efficient access to genealogies and calculation of genetic statistics over the data. We apply this technique to human data from the 1000 Genomes Project, Simons Genome Diversity Project and UK Biobank, showing that the genealogies we estimate are both rich in biological signal and efficient to process.

## Introduction

Using a tree to encode evolutionary relationships is a fundamental organising principle in biology. From Darwin’s speculative sketches [Darwin, 1987] and Haeckel’s phylogenetic imagery [Haeckel, 1866] to modern syntheses encompassing all species of life [Hinchliff et al., 2015], trees elegantly encode and summarise the outcomes of evolutionary processes. Many different methods now exist to infer these evolutionary trees from real-world data [Felsenstein, 2004] and such trees have many applications [Yang and Rannala, 2012]. However, a tree can only be used to describe the ancestry of a set of DNA sequences if these sequences are transmitted as a single unit down the generations. Anything that causes different parts of a sequence to come from different ancestors results in a history that cannot be described by a single tree, but instead requires a network [Morrison, 2016]. This presents difficulties when inferring ancestry within a sexually reproducing species, where DNA is inherited from both mother and father through recombination.

The need for structures more general than trees to describe ancestry has long been understood [Ragan, 2009]. Many different approaches to such “phylogenetic networks” exist, modelling the non-vertical transmission of genetic information such as horizontal gene transfer and hybridisation [Morrison, 2016]. The Ancestral Recombination Graph, or ARG [Griffiths, 1991, Griffiths and Marjoram, 1996], models the network arising from inheritance in sexually reproducing species, encoding the recombination and common ancestor events that occurred in the history of a sample. In principle, ARGs contain all the information knowable about genetic history, and are widely acknowledged as being centrally important to population genetics [Minichiello and Durbin, 2006, Arenas, 2013, Gusfield, 2014, Rasmussen et al., 2014]. However, practical applications have been limited by the fact that inferring ARGs from real data is a prohibitively expensive computational problem. Finding an ARG with the minimum number of recombination events required to explain a set of input sequences is NP hard [Bordewich and Semple, 2005, Wang et al., 2001], and the scope of methods computing such ‘MinARGs’ is therefore severely limited [Hein, 1990, Song and Hein, 2005]. Without this minimality criterion, there exist both polynomial time algorithms [Gusfield et al., 2004, 2007] and methods using various techniques to reduce the state space [Griffiths and Marjoram, 1996, Kuhner et al., 2000, Fearnhead and Donnelly, 2001], but in practice these are too slow to apply to even moderately sized data sets. Several heuristic methods have been explored [Song et al., 2005, Minichiello and Durbin, 2006, Parida et al., 2008, O’Fallon, 2013, Mirzaei and Wu, 2016] but most are limited to tens of samples and a few thousand variant sites. The `ARGweaver` program [Rasmussen et al., 2014] is the current state-of-the-art and a substantial advance over earlier methods, as it performs statistically rigorous inference of ARGs over tens of thousands of variant sites. However, computational time grows very quickly with the number of input samples, and anything more than a few tens of sequences is infeasible. The widespread use of ARGs is also hindered by the lack of a standard means of interchange and toolkits to process such data. Despite several efforts to standardise [Cardona et al., 2008, McGill et al., 2013], adoption remains practically non-existent. Thus, the status of the ARG remains as it has been for many years: a structure that is widely agreed to be fundamental to our understanding of the ancestry of biological populations, but one which is practically never used.

Here we introduce a method, `tsinfer`, that removes these barriers to the adoption of ARGs in the analysis of genetic variation data. Crucially, `tsinfer` vastly expands the scale over which ancestry can be inferred, simultaneously increasing the number of variant sites and sample genomes that can be analysed by several orders of magnitude. In a simulation study, we show that this enormous increase in computational efficiency is not at the cost of accuracy: `tsinfer` infers ancestry with fidelity comparable to the state-of-the-art. Moreover, we show that the data structure produced by `tsinfer`, the *succinct tree sequence* (or tree sequence, for brevity) [Kelleher et al., 2016, 2018] has the potential to store genetic data for entire populations using a fraction of the space that would be required by present-day methods. As an encoding of the data directly based on the evolutionary history of the samples, many statistics can be computed efficiently using this structure—indeed, this efficiency is key to the scalability of `tsinfer` itself. The tree sequence toolkit (or `tskit`) is a free and open source library providing convenient access to these efficient algorithms. Thus, the two main practical obstacles to using ARGs (the lack of efficient inference methods and software to process the output) have been removed. We demonstrate this utility by applying `tsinfer` to three large-scale human data sets (1000 Genomes [1000 Genomes Project Consortium, 2015], the Simons Genomes Diversity Project [Mallick et al., 2016] and UK Biobank [Bycroft et al., 2018]) and show how biological signals can be easily inferred from the resulting genealogical representation.

## Results

### Succinct tree sequences

The tangled web of ancestry describing the genetic history of recombining organisms is conventionally encoded via common ancestor and recombination events in an ARG. An equivalent (and arguably simpler) way of viewing this process is to regard the ancestry of the sample as a sequence of marginal trees, each encoding the genealogy for a particular segment of DNA [Hein, 1990]. As we move along a chromosome, recombination events in the history of the sample alter the trees in a well-defined way [Song and Hein, 2005], and adjacent trees tend to be highly correlated. The *succinct tree sequence* is a recently introduced encoding for recombinant ancestry that takes advantage of the correlations between adjacent trees [Kelleher et al., 2016, 2018]. This is done by recording edges that are shared by multiple adjacent trees once, rather than storing each tree independently. This simple device explicitly captures the shared structure among trees and leads to very efficient algorithms for processing them [Kelleher et al., 2016]. Although essentially equivalent, we avoid describing the succinct tree sequence as an ARG because there is an important distinction between the two structures: an ARG encodes the *events* that occurred in the history of a sample, whereas a tree sequence encodes the *outcome* of those events. In particular, recombination events are not explicitly recorded in a tree sequence.

One benefit of explicitly capturing the shared tree structure inherent in recombinant ancestry is the potential for a dramatic reduction in the space required to store genetic variation data. Such information is usually encoded as a matrix in which the columns correspond to samples (a human individual, for example) and the rows correspond to sites along the genome at which variation is observed (Fig 1A). If we have n samples and m sites we need *O*(*nm*) space to store the entire matrix. Studies such as the UK Biobank [Bycroft et al., 2018] already have data for hundreds of thousands of samples, and such large sample sizes are expected to become increasingly common [Stephens et al., 2015]. As many species have millions of variant sites per chromosome, storing and processing such huge matrices is a serious burden. Tree sequences provide an elegant solution. The variation that we observe in present-day samples is the result of mutations that occurred in the past, in the ancestors of our samples. If we know the genealogy at a particular variant site we can fully describe the variation at that site by recording the presence of these mutations in the ancestors in question (Fig. 1B). Mutations at a given site are typically rare, so we can encode the observed variation among millions of samples by storing the genealogy together with a relatively small number of mutations. The resulting savings in storage space are dramatic. Fig. 1C shows the space required to store variation data for simulations of up to 10 million human-like chromosomes, extrapolated out to 10 billion. In this idealised case, storing the genotype data in the most widely used format (VCF) would require 23PiB (i.e., approximately 23,000 1TiB hard disks) whereas the tree sequence encoding would require only around 1TiB. Thus, if we were able to store variation data using the tree sequence encoding we could store and process any conceivable data set on a present-day laptop.

**Figure 1.**
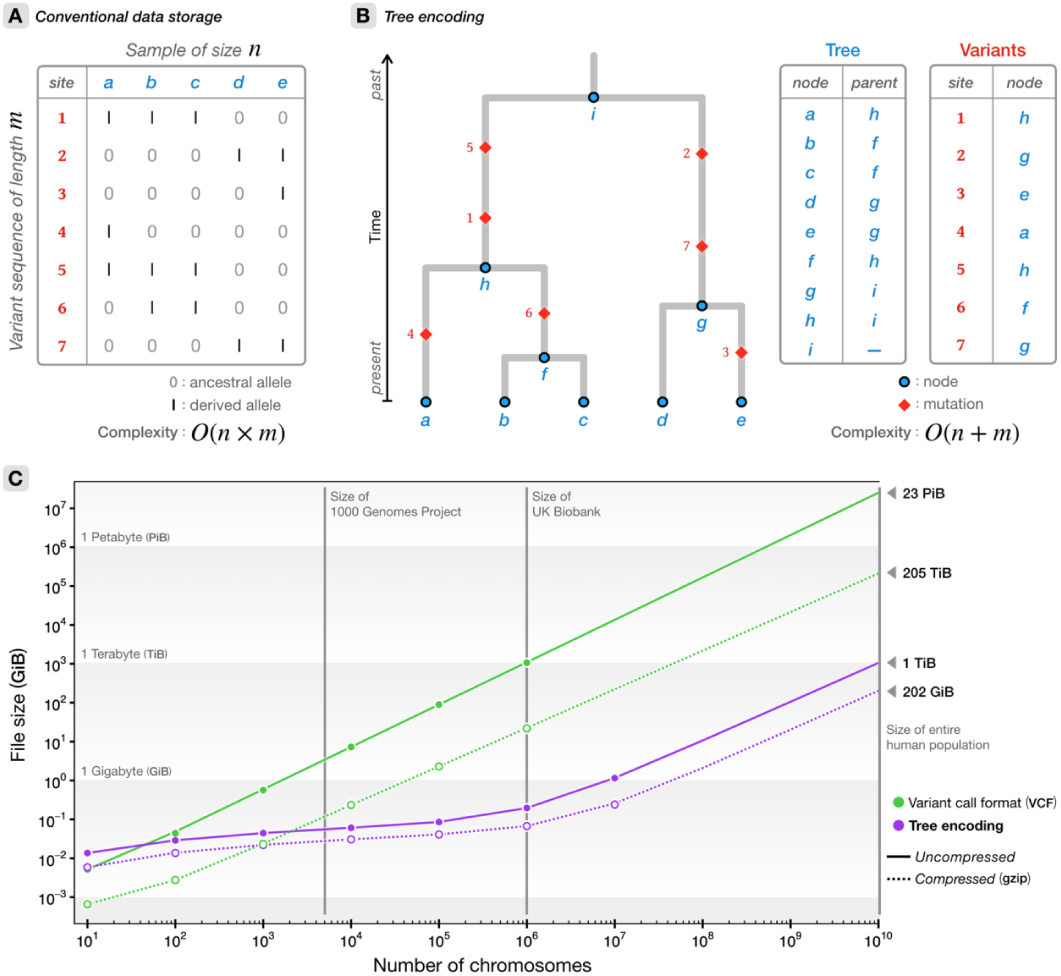
Comparison of tree sequences with standard methods for storing genetic variation data. A). The variant matrix which underlies conventional storage methods for genetic variation data. B). A genealogical encoding for data; if we know the tree we can store each variant site in constant space. C). Estimated sizes of files required to store the genetic variation data for a simulated human-like chromosome (100 megabases) for up to 10 billion haploid (5 billion diploid) samples. Simulations were run for 10^1^ up to 10^7^ haplotypes using `msprime` [Kelleher et al., 2016], and the sizes of the resulting files plotted (points). In each case we show the original tree sequence file uncompressed and compressed. We also show the corresponding variation data encoded in the standard VCF [Danecek et al., 2011] and compressed using gzip. The VCF files for 10^7^ samples were too large and time-consuming to process. The projected file sizes for VCF/compressed VCF are based on fitting a simple exponential model. Projected files sizes for tree sequences are based on fitting a model based on the theoretical growth of tree sequences [Kelleher et al., 2016]. In all cases, the largest data point was withheld from fitting.

Converting data into a highly compressed form usually requires costly decompression before use. A great advantage of the tree sequence encoding is that we can compute many statistics directly from the trees without decoding the genotypes. For example, computing the frequency of specific variants within subsets is a key building-block of many genetic statistics, and in particular is a fundamental operation when performing genome-wide association studies [Manolio, 2013]. The algorithm for recovering trees from the encoded tree sequence representation allows us to compute such allele frequencies far more efficiently than is possible when working with a raw matrix representation of the data [Kelleher et al., 2016]. Take, for example, the largest tree sequence simulated in Figure 1, which represents a simulated history of 10^7^ chromosomes of 100Mb each. It takes about 2.2 seconds to load this 1.2GiB tree sequence; about 7.5 seconds to iterate over all 650K trees; and about 17 seconds to compute allele frequencies within an arbitrary subset of 10^6^ samples at all 670K sites. In contrast, just decompressing and decoding the corresponding 6.1 TiB of genotype data from BCF (a more efficient binary compressed encoding of VCF) would require an estimated 1.8 hours (based on extracting the first 10K variants using cyvcf2 [Pedersen and Quinlan, 2017]). Moreover, BCF and other existing formats for storing variant data do not consider ancestry in any way. If we wished to store the actual trees from this simulation of 10^7^ samples using the most efficient and popular interchange format (Newick), we would need approximately 256 TiB of space and it would take an estimated 5.3 CPU years to parse (based on BioPython’s [Cock et al., 2009] Newick parser—one of the most efficient available—taking about 262 seconds per tree).

### Inference algorithm

DNA sequences can be considered mosaics of sequence fragments that have been inherited from recent ancestors via an error-prone copying process. Likewise, these recent ancestors are themselves mosaics, copied imperfectly from yet older ancestors. As we go further back in time these ancestral sequence fragments (or haplotypes) become shorter, as recombination randomly breaks up the contributions of different ancestors over the generations. Our method is based on the premise that if these ancestral haplotypes were known, it would be possible to infer a plausible copying history for large numbers of input DNA sequences. Critically, this approach means that we do not need to compare sample haplotypes with each other, and we avoid the quadratic time complexity that this implies (which for large data sets, is insurmountable).

We do not usually know these ancestral haplotypes, but we can attempt to infer them. If we assume that the contemporary variation we observe at each site on the genome is the result of a single mutation, then we know that every sample haplotype that has the mutated (or derived) state at this site must have inherited it from a single ancestor. Moreover, these samples will also have inherited some fragment of the ancestral haplotype *around* the focal site. For mutations that recently arose, this shared haplotype will tend to be long, as recombination will not have had time to break it up. Conversely, for ancient mutations the shared ancestral haplotype will tend to be rather short (Fig. 2).

**Figure 2.**
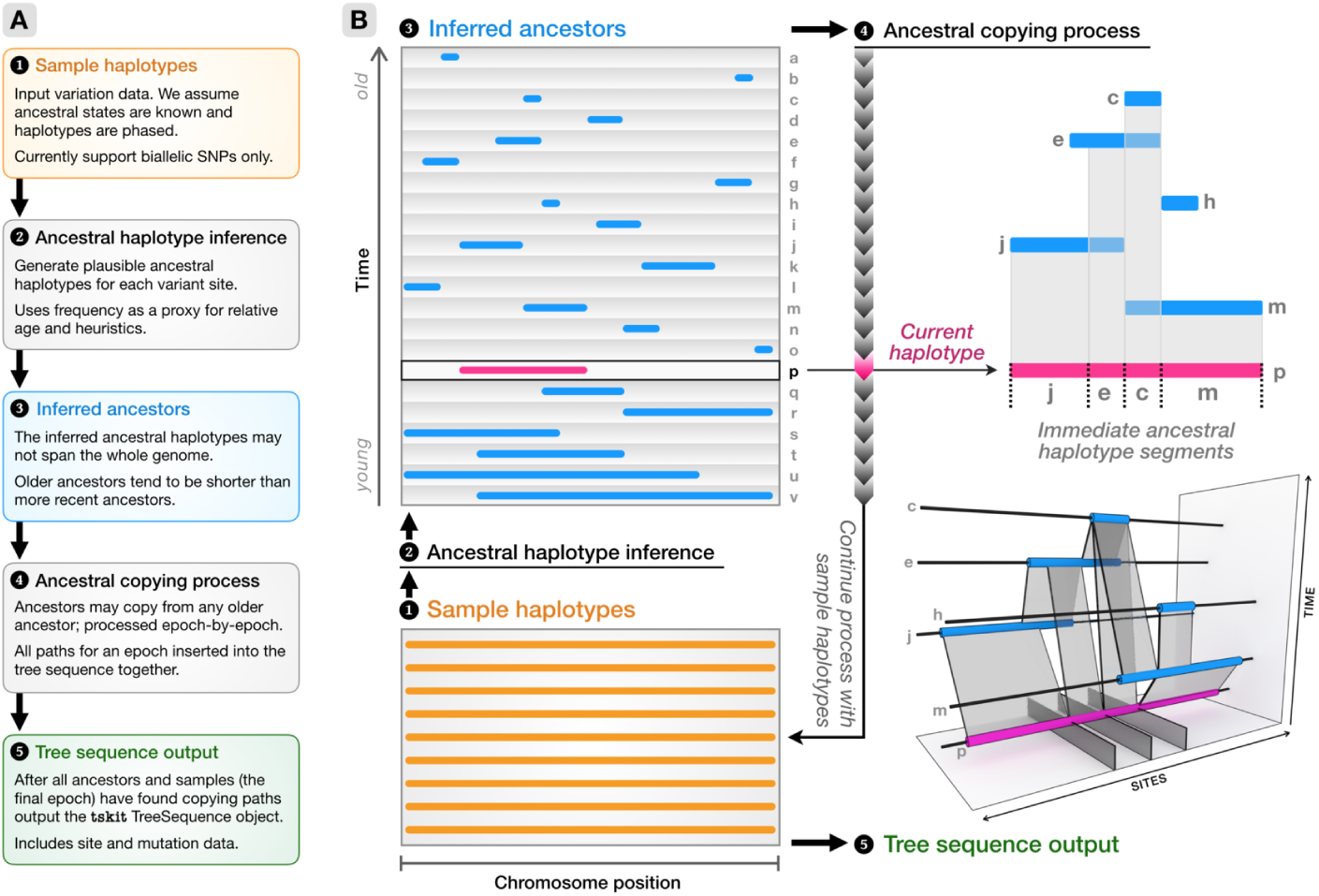
A schematic of the major steps of the inference algorithm. Starting from a set of sample haplotypes extending over the genome (1), we use the ancestral haplotype inference method (2) to reconstruct fragments of ancestral sequence (3), then infer copying paths among these ancestors (4). The ancestral copying process is shown on the right, using an arbitrary haplotype (*p*) for illustration. As we move from left to right along *p* we infer that it has most recently copied from *j*, *e*, *c* and then *m*. Incorporating the copying history of all older haplotypes (for example, *m* copied partly from *c* and partly from *h*), partial coalescent trees emerge in the bottom-right panel. Once copying paths have been found for all ancestors and samples, we output a `tskit` tree sequence(5).

The first step in our algorithm is to generate ancestral haplotypes based on the variation present in the sample sequences. The approach that we use to estimate ancestors is a simple heuristic, which we describe in detail in the Methods. Briefly, we first use the frequency of the derived state at each site as an approximation of the relative age of the corresponding ancestor (we assume that the ancestral and derived states are known). Then, for each ancestor we work outwards from the focal site taking a consensus value among samples carrying the derived state at the focal site. Although heuristic, this method is reasonably accurate and robust to errors (Fig. S3). Furthermore, the approach for generating ancestors is independent of later steps in the algorithm and improved methods can be developed.

After we have estimated ancestral haplotypes, we must then infer how they relate to each other. We do this using a variation of the Li and Stephens (LS) model [Li and Stephens, 2003], one of the most important techniques in contemporary large-scale genomics [Lunter, 2018]. The LS model regards a haplotype as an imperfect mosaic of the haplotypes in a reference panel, and is defined using a Hidden Markov Model (HMM). The most likely path for a given haplotype through the reference panel is found using standard HMM algorithms, and the time required to find such paths scales linearly with the number of haplotypes in the reference panel. For a given ancestral haplotype, our reference panel consists of all older ancestral haplotypes. Because our reference haplotypes are ancestral rather than contemporary, we make a slight modification to the standard LS process: alongside the usual 0/1 states, a third haplotypic state is used to represent non-ancestral material from which copying can never occur. Computing the most likely path under the LS model allows us to estimate the immediate ancestor for each segment of DNA in the focal haplotype. Figure 2B shows an example of such a copying path for a focal haplotype and how it copies from different ancestors along its length. Once we have found copying paths for all of our ancestors and all of the input sample haplotypes, we are guaranteed to have complete genealogical trees for every position along the genome, albeit ones that may contain nonbinary nodes (“polytomies”). Furthermore, these copying paths map directly to ‘edges’ in the tree sequence formulation [Kelleher et al., 2018], and so the copying process directly generates a succinct tree sequence. The output tree sequence may optionally be “simplified” [Kelleher et al., 2018] to remove any generated ancestral segments unreachable from the sample nodes. Representing the ancestral haplotypes as a tree sequence lends significant flexibility, as it allows us to combine information from diverse sources; for example, we can use a tree sequence estimated from one data set as ancestors for another (see Applications).

The correspondence between the output of the copying process and a tree sequence is also critical to scalability. Because each non-singleton input site usually corresponds to a single ancestor, the reference panel we match against may contain millions of ancestral haplotypes. Furthermore, this reference panel must be dynamically updated to include more and more ancestors, as our strict time ordering requirements result in haplotypes being able to copy from all older haplotypes. Such requirements are far beyond existing methods for finding likely paths under the LS model, which either require time that is linear in the reference panel size or a linear time preprocessing step [Lunter, 2018]. The key technical advance that makes our method feasible for large samples is an exact solution of the LS HMM that uses the partially built genealogies to greatly speed up calculations (see the Methods for details).

### Algorithm evaluation

We evaluate `tsinfer` for accuracy and scalability using simulated data. We perform population genetic simulations which output a tree sequence as a .trees file. These encode both the simulated genealogies, which we use as a ground-truth, and sample haplotypes, which we use for inference. We compare `tsinfer` to three other tools for ancestral inference. `ARGweaver` [Rasmussen et al., 2014] is the most statistically rigorous, and is considered state-of-the-art. Rent+ [Mirzaei and Wu, 2016] is a heuristic method that is considerably faster than `ARGweaver`. Finally, fastARG (https://github.com/lh3/fastARG) is an unpublished method using an approach similar to Margarita [Minichiello and Durbin, 2006]; we include fastARG in this analysis because it is by far the most scalable method currently available. A degree of error is inevitable in DNA sequence data and methods must be reasonably robust to be relevant to empirical data. We therefore impose a genotyping error process derived from an empirical analysis [Albers and McVean, 2018] on the simulated haplotypes (leading to an observed error of around 0.35%), and assess the performance of tools with and without the presence of these simulated errors.

We evaluate the accuracy of the different methods by comparing estimated tree topologies with the original simulated trees. We use the Kendall-Colijn tree distance metric [Kendall and Colijn, 2016], as it is more sensitive than alternative metrics (Fig. S5) and is robust to the presence of non-binary nodes in trees (see Methods for details). As `tsinfer` does not currently try to estimate node times, we only consider topological similarity and disregard branch lengths. Fig. 3 compares the accuracy of `tsinfer` against other tools on simulated data, with and without simulated genotyping error, for a variety of different mutation to recombination rate ratios. As we increase the mutation rate the accuracy of the inferred trees increases for all tools, because more mutations reveal more information about the underlying tree topologies (we generate mutations under the infinitely-many-sites model and so cannot have recurrent mutations). In this evaluation `tsinfer` is substantially more accurate than `Rent+` and `fastARG`, both with and without error. We can also see that `tsinfer` produces more accurate topologies than `ARGweaver` when the mutation rate is lower than the recombination rate, and has comparable accuracy with higher mutation rates; however, `ARGweaver` is somewhat more robust to the presence of errors than `tsinfer`. Similar results are obtained when we simulate more complex demography in Fig. S6. In contrast, Fig. S7 shows that `tsinfer`’s inference accuracy is substantially higher than `ARGweaver`’s in the presence of a selective sweep, suggesting that our nonparametric approach is more robust to such departures from the assumptions of the coalescent model.

**Figure 3.**
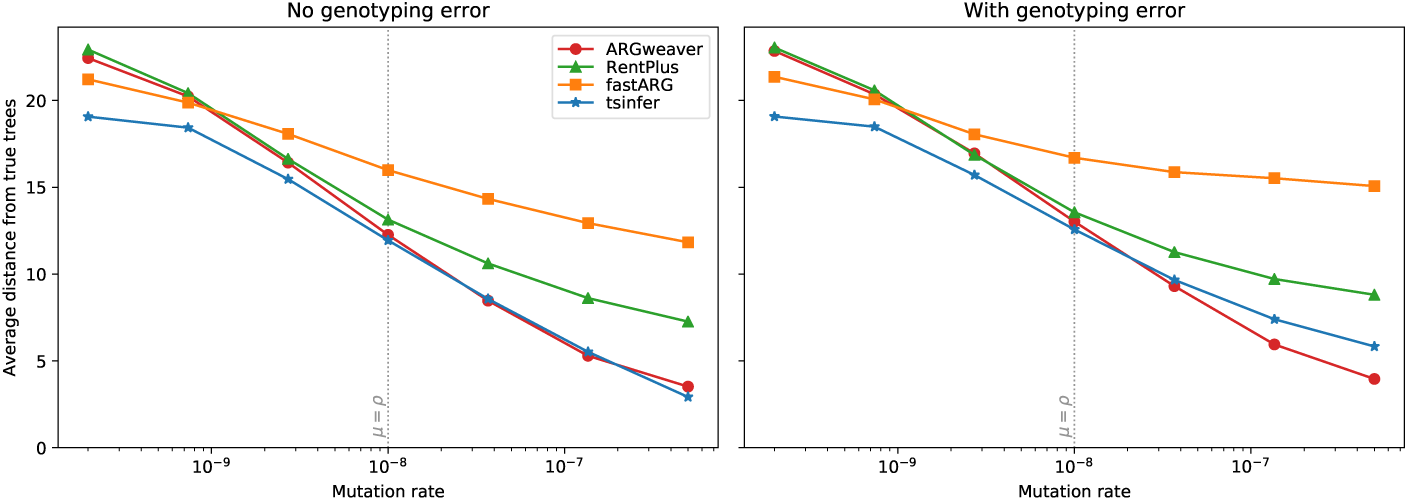
Accuracy of ancestry inference under different methods (lower values indicate greater accuracy). Coalescent simulations for 16 sample haplotypes of 1Mb in length under human-like parameters (*N_e_* = 5000, with recombination rate *ρ* = 10^−8^ per base per generation) and an infinite sites model of mutation were simulated using `msprime` [Kelleher et al., 2016]. The reported tree topology distance is the Kendall-Colijn metric, weighted by the genomic distance spanned by each tree. Each point is the average of 100 independent replicates at a given mutation rate. The point where mutation rate equals recombination rate (similar to humans) is marked with a vertical dotted line. Standard errors are smaller than the plotted symbols in all cases.

It is notable how small the data sets used to compare these tools are: we are using 8 diploid samples, and, where the mutation equals the recombination rate, an average of 662 variant sites. This is driven by necessity as `ARGweaver` is very CPU intensive. Supplementary Fig. S9 shows how the CPU time scales for these tools for up to 100 samples, and illustrates the vast differences in processing time required: `ARGweaver` requires hours while `fastARG` and `tsinfer` require fractions of a second. `Rent+` is substantially faster than `ARGweaver`, but is still orders of magnitude slower than `fastARG` and `tsinfer`. Although `fastARG` is slightly faster than `tsinfer` for tiny data sets, Fig. S10 shows that `tsinfer` is far more efficient for inference involving tens of thousands of mammalian-scale chromosomes.

Succinct tree sequences have great potential as a means of storing genetic variation data, as simulated tree sequence files are many times smaller than the corresponding compressed genotype matrices for large sample sizes (Fig. 1). To evaluate `tsinfer`’s compression performance on large-scale data we inferred tree sequences from simulated sequence data and compared the size of the inferred .trees files to the equivalent compressed VCF (Fig. S11). While compression performance varies with the ratio of mutation to recombination rate, in our example of 100K samples, sequences greater then 50Mb in length have inferred tree sequence files ranging from 10-20 times smaller than the corresponding compressed VCF. Fig. S12 shows that in many cases `tsinfer` can estimate large-scale tree sequences that are even smaller that the original simulated files.

### Applications

To evaluate `tsinfer`’s performance on empirical data we constructed tree sequences for three data sets on human chromosome 20: the 1000 Genomes Project (TGP), consisting of low-coverage whole genome sequencing data from 2504 individuals across 26 worldwide populations [1000 Genomes Project Consortium, 2015]; the Simons Genome Diversity Project (SGDP), consisting of high coverage sequencing data from 278 individuals from 142 worldwide populations [Mallick et al., 2016]; and the UK Biobank (UKB), consisting of SNP array data from 487,327 individuals within the UK [Bycroft et al., 2018]. Table 1 summarises input data, inferred tree sequences and computing resources required. For UKB, we considered multiple strategies, augmenting the data with ancestors inferred from TGP and subsets of haplotypes from the UKB itself as potential ancestors. For each data set we used statistically inferred haplotypes as input.

**Table 1.** Summary of input data, output tree sequences and computing resources required for TGP, SGDP and UKB chromosome 20. Input sizes reported are of `tsinfer`’s input .samples files, which uses the Zarr library (https://zarr.readthedocs.io/) to achieve similar compression levels to BCF. File sizes are reported using binary multipliers (i.e., 1M = 2^20^ bytes); all other values use decimal multipliers (i.e., 1M = 10^6^). The times reported are the total wall clock time required to produce the output tree sequence from the `.samples` file on a server with two Xeon Gold 6148 CPUs (40 cores in total; no hyperthreading) and 187GiB of RAM. For SGDP, TGP and UKB we used the standard `tsinfer` inference pipeline. In UKB+TGP, we matched the UKB samples to the inferred TGP tree sequence (time reported is just for sample matching phase). In UKB+UKB we incrementally added samples from UKB to the ancestors inferred from UKB (see text).

|  | Input |  |  | Output |  |  |  | Resources |  |
| --- | --- | --- | --- | --- | --- | --- | --- | --- | --- |
| | $n$ | sites | size | nodes | edges | trees | size | time | RAM |
| SGDP | 277 | 348K | 15M | 236K | 1.7M | 196K | 83M | 5m | 3.6G |
| TGP | 2504 | 860K | 135M | 735K | 7.3M | 550K | 296M | 2h | 11G |
| UKB | 487K | 15.8K | 1.6G | 1.9M | 484M | 15.8K | 14.5G | 3h | 160G |
| UKB+TGP | 487K | 15.6K | 1.6G | 5.5M | 185M | 15.6K | 5.8G | 15h | 66G |
| UKB+UKB | 487K | 15.8K | 1.6G | 2.0M | 62M | 15.8K | 2.1G | 50h | 40G |

Across chromosome 20, the TGP data consisted of 860K sites after filtering (see Methods for details). After inferring ancestors and matching sample haplotypes to these ancestors, we obtain a 297MiB tree sequence (109MiB compressed, compared to the 141MiB BCF encoding of the same genotypes). Loading the tree sequence required c. 3 seconds; iterating over all 550K trees c. 0.6 seconds; and decoding all genotypes c. 9 seconds. In comparison, decoding the same genotypes from BCF required c. 15 seconds using cyvcf2. The SGDP data consisted of 348K sites after filtering, and this resulted in an 83MiB tree sequence (28MiB compressed, compared to 11MiB BCF encoding of the same genotypes). Loading the tree sequence required c. 1.6 seconds; iterating over all 196K trees c. 0.1 seconds; and decoding all genotypes c. 1.8 seconds. These results demonstrate the feasibility of representing existing data sets through tree sequences, with file sizes comparable to current standards and excellent analytical accessibility.

To assess the validity of the inferred tree sequences we computed a series of metrics summarising reconstructed ancestral relationships. We first calculated the number of edges for each sample, which measures the extent to which an individual’s genome can be compressed against the inferred ancestors. In TGP, samples have an average of 648 edges (with a median length of 44kb and an average N50 of 236kb), with those of African ancestry having a greater number (750) than those of European (551) or Asian (665) ancestry (Fig. S13A and Fig. S14). These findings are likely to primarily reflect known differences in the long-term effective population size, though will also be affected by sampling strategy and error modes. We find higher values in SGDP reflecting the lower sample sizes (overall average: 1113 edges, African ancestry: 2178, European ancestry: 803 and Asian ancestry: 879; Fig. S13B and Fig. S15). In both data sets we identified a few outlier samples with very high edge counts, suggesting error (see Methods).

We next considered whether the inferred tree sequences could be used to characterise ancestral relationships in TGP and SGDP by computing, for each individual, the population distribution of their genealogical nearest neighbours

(GNN). Given *K* sets of reference nodes (e.g., the samples for each of the 26 TGP populations), the GNN statistic for a specific node is a *K*-vector describing the proportion of its immediate neighbours within the tree from each of these reference sets (see Methods). We find that, despite the noise generated by uncertainty in tree reconstruction (manifest as polytomies), the chance nature of the genealogical process and data error, the tree sequences can characterise global population structure (Fig. 4A, B), within-population relatedness (Fig. 4C) and identify regions of differential ancestry within an individual (Fig. 4D). These analyses demonstrate the potential of interrogating the inferred genealogical structure at different resolutions to describe both broad and fine-scale patterns in contemporary human genomic diversity. The statistics involved can be calculated very efficiently: using 8 threads, computing GNNs for every sample required c. 16 and 30s, respectively, for the SGDP and TGP data sets (Xeon Gold 6148 CPU).

**Figure 4.**
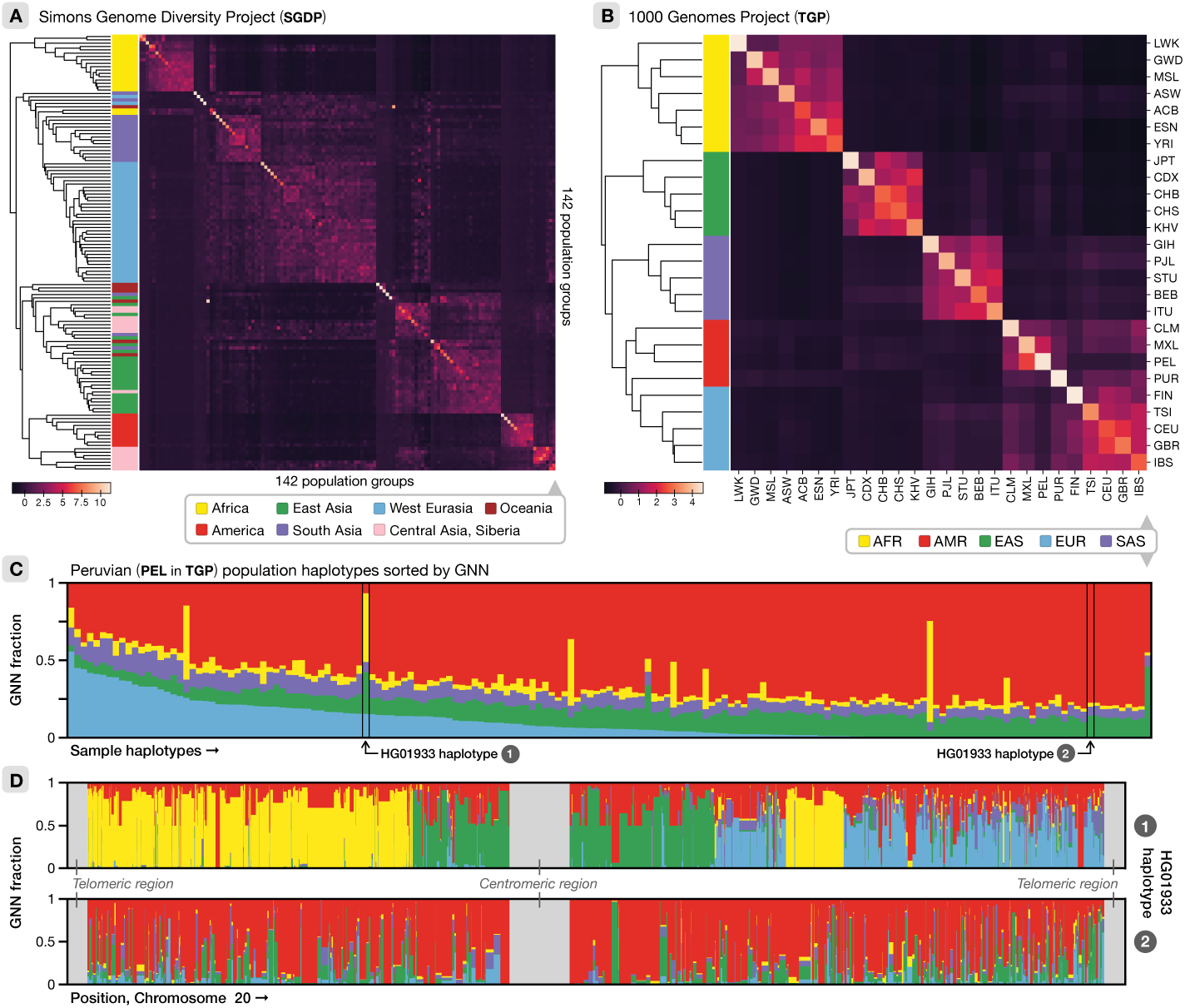
Tree sequence characterisation of global genome diversity. A). Z-score normalised GNN proportions for SGDP by population. The GNN matrix was first z-score normalised by column and the rows then hierarchically clustered. See Fig. S16 for a larger version with population labels. B). As for (A), but for the TGP data. C). Average GNN proportions for all individuals within the PEL population in TGP. Colours indicate continental-level groupings. D). The approximate GNN proportions across the chromosome for the two haplotypes of HG01933, from the PEL population in TGP. HG01933 was chosen as an example of an individual showing evidence of very recent admixture.

Finally, to assess the performance of `tsinfer` on vast data sets we analysed the ∼500K individuals within UK Biobank. The sparsity of variant sites and inherent lack of rare variants in the UKB SNP array data is insufficient for accurate ancestor inference directly. However, we considered two alternative strategies: using ancestors estimated from other data and using subsets of the sample to act as proxies for ancestors. In the first approach, we matched the UKB haplotypes to the tree sequence inferred for the TGP, generating a 5.8GiB tree sequence (2.1GiB compressed). Fig 5A shows the self-reported ancestry in UKB tallies with TGP GNN values and adds granularity. Furthermore, by analysing the copying patterns in this tree sequence, we found 9 individuals that are likely to be in both the TGP and UKB data sets (see Methods).

**Figure 5.**
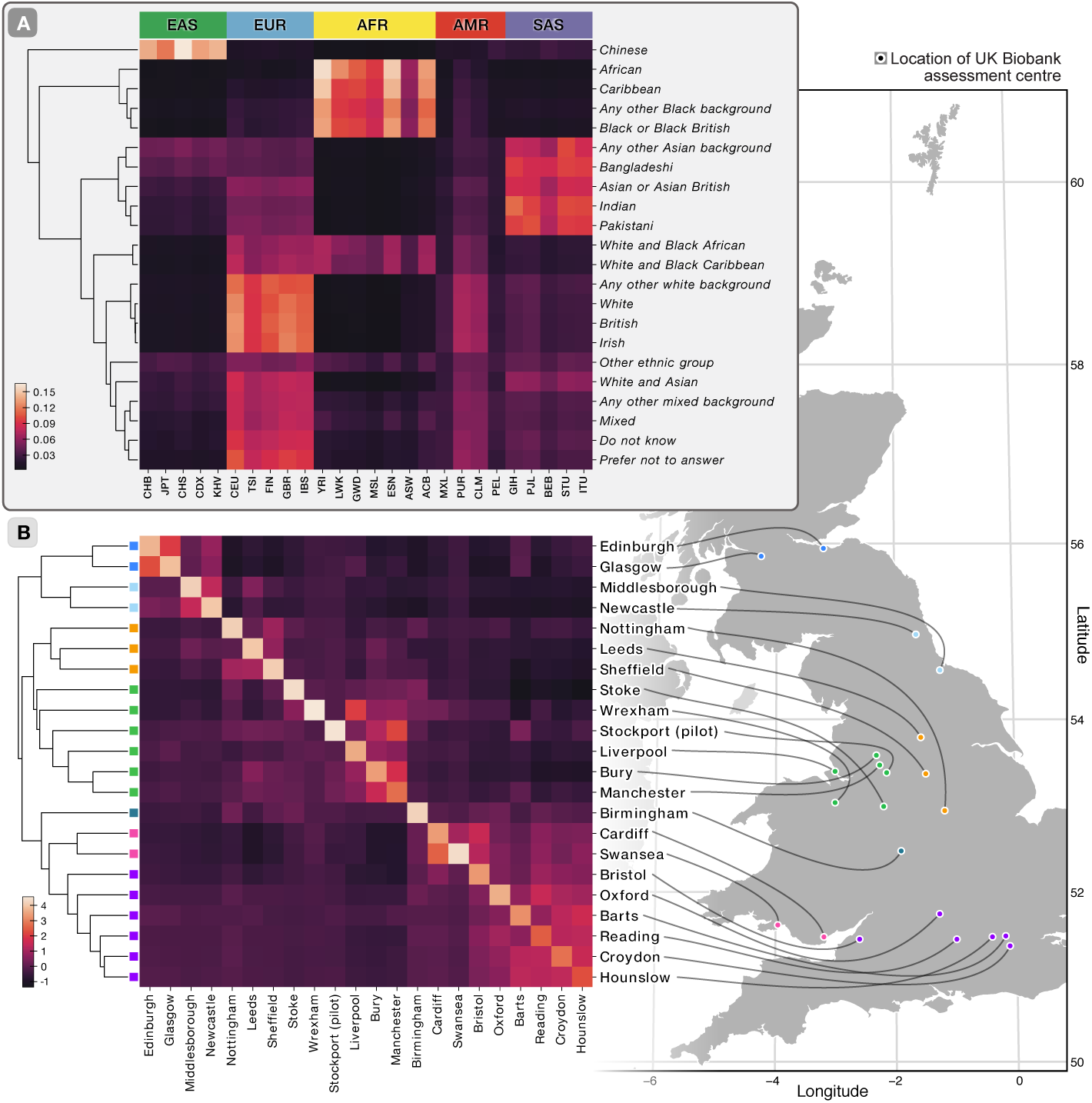
Tree sequence characterisation of the UK Biobank data. A). Ancestral relationships captured from GNN proportions in the tree sequence constructed from TGP data, relative to TGP populations, organised by self-reported ancestry. Rows are clustered hierarchically. B). Z-score normalised GNN proportions in the tree sequence constructed by iteratively augmenting UKB derived ancestors with sample haplotypes for 629,246 haplotypes self-reported as of British ancestry (the 262,142 sample haplotypes used for ancestor augmentation were removed from this analysis). The GNN proportions here were calculated relative to sample enrolment centre, i.e., reporting the fraction of each sample’s genealogical nearest neighbours that enrolled at a given centre. The columns in the GNN matrix were z-score normalised before the rows were hierarchically clustered.

Our second strategy for improving ancestor inference involves sequentially adding subsets of the sample itself as potential ancestors, which we deployed to investigate structure within the UKB. By iteratively adding more and more samples, our method of generating putative ancestors from shared recombination breakpoints (“path compression”; see Methods) is able to construct many more ancestors that would be possible if we added all of the extra samples at once. Thus, we began by updating the ancestor’s tree sequence with the paths for two arbitrarily chosen samples; then updating the resulting tree sequence with the paths taken for four other samples; and then again for eight; and so on up to 131,072. After matching all 1M sample haplotypes against these augmented ancestors we obtained a 2.1GiB tree sequence (928MiB compressed, compared to the equivalent 1.4GiB BCF file). Loading this tree sequence required c. 9 seconds and iterating over all 15.8K trees c. 11 seconds. Decoding genotypes for the first 1000 sites required 9.5 seconds; in comparison, decoding the genotypes for the first 1000 sites in the original BGEN file using the bgen C library required 49 seconds. Analysis of the GNN structure of the tree sequence demonstrates strong geographical clustering of relatedness at this level of resolution, with connections between enrolment centres reflecting geographical proximity (for example connections between centres within Scotland and Wales; Fig. 5B). Although signals of population structure are evident here, further work is required to understand the implications for statistical analysis of association.

## Discussion

Inferring genealogical relationships among individuals from patterns of genomic variation is a long-standing problem in evolutionary biology that connects to many of the fundamental forces and events that shape a species, including mutation, recombination, demographic processes and natural selection. However, our ability to infer such histories has, to date, been limited by the computational complexity of the problem. The work presented here represents a major advance by providing a principled, yet scalable approach that can be applied to data sets of unprecedented size. While the algorithms presented are both heuristic and deterministic, the approach of breaking down the problem into inferring relative variant age, ancestral haplotypes and the genealogical relationships between these ancestors results in a modular framework that scales to vast sample sizes (Fig. S10). Moreover, each component can be improved independently, even potentially accommodating uncertainty through stochastic approaches.

Nevertheless, the method does make a number of fundamental assumptions. First, we assume that each variant in a population has a single mutational origin. While this is unlikely to be true in practice (particularly in large samples), our ancestor estimation method is likely to find the dominant ancestral haplotype. Recurrent and back mutations will therefore not be handled well by the current algorithm, though in principle could be addressed by iterative approaches. Second, we assume that frequency is a proxy for relative variant age. Importantly, our algorithms only require accuracy about relative age within genealogically connected parts of the tree sequence. Under simple demographic models, we estimate that relative frequency indicates relative age for roughly 90% of closely located pairs of variants (Fig. S1). In theory, methods for dating genomic variants could be used to improve ancestor estimation and also assign dates to nodes within the tree sequence, though these remain open problems. Third, we assume that the ratio of mutation to recombination is sufficiently high to use mutations as the starting point for ancestor inference. However, the path compression approach used here (see Methods) essentially identifies additional ancestors through shared recombination events and, within the SNP array analysis on UKB, has been shown to perform well, compensating for the low variant density and lack of rare variants. Finally, the current methodology works well for low error rates, but its performance is degraded by genotyping and, in particular, haplotype phasing error. In the future, population-scale high coverage and routine long-read genome sequencing will reduce the source of such errors and it may be feasible to construct steps of the algorithm that are more robust than those currently implemented.

Tree sequences have multiple potential applications. The most obvious is as an efficient lossless storage format for population-scale data sets. While compression performance on simulated data (Fig. S11) is close to the theoretical possibilities shown in Fig. 1, `tsinfer`’s compression of real data does not currently fulfil this great potential. More careful modelling of the complexities of genetic data—in particular the various error modes—will likely be required to effectively compress millions of whole genomes. Nonetheless, compression performance is comparable to existing formats while providing exceptional analytical accessibility. The integration of genealogical relationships with genomic variation data has value beyond population and personal genetics, for example in potentially correcting for the differential geographical confounding of rare and common variants in genetic association. However, the combined analysis of the UKB and TGP data sets demonstrates the potential of also using tree sequences to integrate data sources and, more generally, to build a reference tree sequence structure for human genomic variation that can be updated as new variants are discovered. Such a structure, coupled with efficient algorithms that make use of the tree structure, such as the LS algorithm deployed here, could enable (and make optimally powerful) diverse statistical genetic operations including genotype refinement, genotype imputation and haplotype phasing. It could also be used to share data effectively and in a manner that preserves privacy, by describing data sets in terms of representation of ancestors rather than individual samples.

## Methods

### Age of alleles

The first step in our algorithm is to estimate the relative time at which the mutation for each variant arose (we are assuming a single origin for each mutation). Classical results in population genetics provide a theoretical expectation for the age of an allele based on its frequency [Kimura and Ota, 1973, Griffiths and Tavare, 1998]. There are several existing methods for estimating allele age, but are either computationally expensive or require detailed knowledge about historical population processes [Ormond et al., 2015, Nakagome et al., 2016, Smith et al., 2018], although a more efficient non-parametric method has recently been introduced [Albers and McVean, 2018].

For our purposes, frequency provides a computationally inexpensive and surprisingly accurate proxy for relative allele age (Fig. S1). This is because we only need the age ordering of alleles to be *locally* accurate: if variants on opposite ends of a chromosome are incorrectly ordered it makes little difference to the outcome of our algorithms, since the ancestral haplotypes involved are unlikely to overlap. Although these estimates could certainly be improved by using the methods mentioned above, our current algorithms for inferring ancestral haplotypes and computing copying paths require a single origin for each mutation. The frequency estimate coupled with our algorithm for generating ancestral haplotypes (see the next subsection) guarantees this property, simplifying the overall process.

### Inferring ancestral haplotypes

Once we have assigned an age order to sites, the next step in our inference process is to generate a set of putative ancestral haplotypes. We assume that there are two alleles at every site: the ‘ancestral state’, which was inherited from the ancestor of the entire population and the ‘derived state’ which is the result of a mutation that occurred on the ancestor of the samples carrying this allele. We assume that these ancestral and derived states have been identified via existing methods (e.g., Keightley and Jackson [2018]). Each variant site is therefore the result of a mutation that occurred on an ancestor: samples that inherit from this ancestor have the derived state, and the rest carry the ancestral state. By definition this ancestor carries the derived state at the site in question; our task then is to reconstruct the state of the ancestor *around* this focal site.

For a given focal site *l* let *S* be the set of samples that carry the derived state. We are attempting to reconstruct the ancestral haplotype *a* on which the mutation occurred, and so we begin by setting *a_l_* = 1, following the usual convention of labelling the ancestral state for a site 0 and the derived state 1. For all other sites 1 ≤ *k* ≤ *m*, *k* ≠ *l* we set *a_k_* = −1, indicating non-ancestral material that cannot be copied from; these non-ancestral values will be overwritten for sites around *l* where we can estimate the state of the ancestor. We then work leftwards and rightwards from *l* independently, computing the state of the ancestor at each site. The set *S* initially contains the samples that we believe descend from the current ancestor (assuming infinite sites and no error), and we use this set to compute a plausible state at other sites. As recombination modifies the tree topology, we update *S* to remove samples that are no longer in the clade induced by the focal site. We stop moving left or right from the focal site when we judge that we no longer have sufficient information to construct the ancestral haplotype. We use heuristics to determine when to remove a particular sample from *S* and when to end the ancestral haplotype.

Figure S2 illustrates a simplified example of this process, showing the ancestral haplotype estimated at the focal site 8. We begin by setting *S* = {*e*, *f*, *g*, *h*}, i.e. the set of samples that carry the derived allele at site 8. We then proceed rightwards, considering each site in turn. For younger sites, the corresponding mutation cannot have occurred yet by definition, and so we always set the ancestor to 0 at these sites (e.g. 9 and 10). When we reach a site that is older than the focal one, we compute a plausible value for our ancestor by taking the consensus among the samples in *S*. For example, at site 11 the estimated value for the ancestor is 1 because all haplotypes in *S* carry 1; similarly, at site 13, three of four samples in S carry 1, and so the consensus is 1 (the consensus can also be 0, as in site 4). We interpret disagreement with the consensus value as evidence that the samples in question have recombined away to another part of the tree. Thus, after we have computed the ancestor’s state at a site remove any samples from S (“evict”) that conflict with this consensus (but see below for a slight modification used in practise). In the example, we therefore evict *h* at site 13 and *g* at site 17. We continue rightwards in this way until we determine that we have insufficient information to accurately estimate ancestors. The heuristic we have chosen is to stop when the size of *S* is ≤ half of its original size. After completing the rightwards scan, we then repeat the process independently leftwards.

Variants with an age equal to the focal mutation are considered to be younger than the focal site (and hence always assigned the ancestral state) except in one special case. If several sites exist with precisely equal distribution of genotypes, we assume that these all arose on a single branch of the tree and that no recombination occurred between these sites. We therefore compute consensus values for older sites between these identical focal sites in the usual way, but we do not update *S* when conflicts occur (assuming these to be caused by error). Once outside of the region enclosed by the identical sites, the process outlined above resumes and we update *S* in the usual way.

Although this method is approximate and heuristic, it generates surprisingly accurate ancestors. Fig. S3 shows a plot of the lengths of the estimated vs true ancestors from simulations, colour-coded by the accuracy of the estimated states. We see that there is a strong bias towards ancestral haplotypes being longer than the truth; this is by design, as long haplotypes can be compensated for by the copying process, but short haplotypes cannot. Inferred haplotypes are also quite accurate, with many ancestors being inferred perfectly. The process is reasonably robust to genotyping errors, but these can create one notable issue for the algorithm illustrated in Figure S2: some generated ancestors are too short, because errors can lead to samples being prematurely evicted from *S*. To add some resilience to this, we include a slight “dampening” to our eviction rule: we remove a sample from *S* only if it disagrees with the consensus at two consecutive older sites.

### Copying process

Given an input haplotype with *m* sites and a reference panel of *n* haplotypes, the most likely path under the Li and Stephens (LS) model is found using the Viterbi algorithm. In the first phase of this process we proceed site-by-site from left to right. At each site, we compute the likelihood that the input haplotype has copied from a particular reference haplotype given their states and the most likely haplotype at the previous site. Once we have reached the last site and we know the most likely reference haplotype at the end of the sequence, we then trace back through the sites, switching to other haplotypes where required. The overall time complexity is therefore *O*(*nm*) to find a copying path for an input haplotype, since we must compare with all *n* reference haplotypes at each of the m sites. In `tsinfer` the reference panel is the set of inferred ancestral haplotypes. Because we may have a different ancestor for every site, n ≈ m, and the time complexity of finding a copying path for an ancestor is therefore *O*(*m*^2^). Modern sequencing data sets contain millions of variant sites and standard LS methods are therefore not feasible.

In `tsinfer` we use the LS model to find a Viterbi path for an input haplotype through the set of ancestors. Each copying path generated is equivalent to a set of edges in a tree sequence [Kelleher et al., 2018] where the child is the focal haplotype and the parents are copied ancestors. Therefore, as we go forwards in time finding copying paths for younger and younger ancestors, we are also incrementally building a tree sequence encoding the state of these ancestors. We use this partially built tree sequence representing a subset of the ancestors as the substrate for computing LS copying paths for subsequent ancestors. The powerful computational properties of the tree sequence data structure allow us to find exact copying paths far more efficiently than is possible using standard methods.

The algorithm for computing Viterbi paths using a tree sequence works in the same way as the standard method. We first proceed from left-to-right, computing a likelihood of copying from each ancestor at every site and recording the locations of potential recombination events. Once we have reached the last site, we trace back as before, resolving a full copying path from the stored information. The difference in our method is that we avoid needing to compute and store a likelihood for each reference haplotype by using the tree sequence to compress the associated likelihoods. Each ancestor corresponds to a node in the marginal tree at a given site, and we compress the likelihoods by marking any node that has the same likelihood as its parent with a special value. The number of distinct likelihoods on the tree is then small, and we can store a list of the nodes that represent whole subtrees. Updating the likelihoods at a given site is then straightforward. We compute the likelihood for each node by reasoning about the state of the input haplotype and the location of the site’s mutation in the tree. Having updated the likelihoods for the nodes corresponding to the compressed subtrees, we can then recompress to take into account the new likelihood values and proceed to the next site. In many cases, moving to the next site will also involve a change in the tree topology and we redistribute the compressed likelihoods accordingly using logic common to other tree sequence algorithms [Kelleher et al., 2016, 2018]. In the interest of simplicity our current implementation does not include a ‘mismatch’ term and only allows for exact haplotype matching. Under these assumptions, we need only five discrete values to encode the node likelihoods in our LS Viterbi algorithm, simplifying the logic considerably. Adding the mismatch term is not difficult (it was present in earlier versions of the algorithm) and we plan to include it in subsequent versions of `tsinfer`. This overview of the LS model on tree sequences is necessarily brief and imprecise, as a full treatment is beyond the scope of this paper. A detailed description and analysis of the generalised model is planned for future work, along with an efficient implementation in `tskit`.

To validate the correctness of our implementation of the copying process, we devised a strong test which we call ‘perfect inference’. In this test we begin with a simulated tree sequence with no mutations and derive the true ancestral segments from it. We then add a specific pattern of mutations that are designed to precisely identify the endpoints of each ancestor and use the resulting ancestral haplotypes as input to `tsinfer`. We then find copying paths for these ancestors and samples in the usual way. Remarkably, using this method we are able to perfectly reproduce the input tree sequence topology, recovering every marginal tree and recombination breakpoint exactly for arbitrarily large inputs. Indeed, the numerical tables [Kelleher et al., 2018] representing the input and output tree sequences are byte-for-byte identical. This is both a strong test for the correctness of our implementation and a reassuring validation of the overall method: if we have accurate ancestors we converge on the true ancestry.

### Path compression

The algorithm for inferring ancestors discussed above is mainly based on the signal arising from mutations, and is only weakly informed by recombination. However, we can also derive information about the ancestors of our sample from recombination events. If we assume that each recombination event is unique, i.e., that all samples that inherit the local haplotype resulting from a breakpoint did so from a single ancestor, we can then estimate the state of this ancestor. Note that this is equivalent to assuming an ‘infinite-sites’ like model for recombinations, an idea with a long history [Fisher, 1954]. We use this signal of shared recombination breakpoints in a specific way, which we refer to as ‘path compression’.

When generating copying paths for successive ancestors, we will often find that subsets of two or more paths are identical. Such identical path subsets is strong evidence for the existence of a single ancestor that consisted of the concatenation of the corresponding haplotype segments. We therefore add this ‘synthetic’ ancestor, and adjust the original identical path subsets to copy from the newly inserted ancestor. Ancestors corresponding to a given allele age are inserted at the same time, and path compression is run at the end of each of these time slices.

Fig. S4 shows a simple example of this process where we compress the shared path subset for the ancestors *g*, *h*, *i*, and *j* into single edges pointing to a new synthetic ancestor. In the top panel we can see (e.g.) that *g* has the copying path (5,12, *c*), (12, 22, *d*), (22, 28, *b*) and *h*, *i* and *j* also contain this path. We therefore create a new ancestor *s* which consists of this path and replace the mappings for *g*, *h*, *i*, and *j* with the single edge (5, 28, *s*). In this way we reduce the overall number of edges required to represent the history, and also provide an extra ancestor for subsequent haplotypes to match against.

### Inference accuracy

We compare the accuracy of `tsinfer` against other inference methods using simulation. We begin by simulating a set of tree sequences which provide the ground-truth topologies. We assess the effect of genotyping errors on inference accuracy by simulating errors on the haplotypes using an empirically determined genotyping error profile [Albers and McVean, 2018]. We then input the corresponding haplotypes (with and without simulated genotyping errors) to the various tools, and measure the difference between the estimated and true tree topologies using tree distance metrics. We repeat this process independently for four tools: `tsinfer`, `ARGweaver`, `Rent+` and `fastARG`. `ARGweaver` requires several parameters to be specified: in all cases we use the known simulation values for the effective population size, mutation and recombination rates. We use the default number of timesteps for time discretisation (20) and for each simulation sample 10 ARGs (one every 20 MCMC cycles) after a burn-in period of 1000 cycles.

Although the succinct tree sequence data structure can fully represent node timing (and hence branch length) information, `tsinfer` does not currently attempt to accurately infer the times of ancestors. Therefore, we limit our investigation of the accuracy of inference to assessing the quality of the inferred topologies, and do not consider timing information in any way. Fig. S5 compares the accuracy of the tools over a number of tree topology distance metrics. For each replicate simulation we compute the average distance between pairs of true and inferred trees along the sequence, weighted by the distance along the sequence that these trees persist. We then report the average distance over the replicate simulations. Metrics are calculated using the R packages `treespace` [Jombart et al., 2017] and `phangorn` [Schliep, 2011]. Some metrics (such as Robinson-Foulds) are undefined for trees that contain non-binary nodes (“polytomies”), which indicate uncertainty in `tsinfer` and therefore occur frequently. To use these metrics in a well-defined way, we also show results where `tsinfer` trees have been randomly resolved into fully bifurcating trees (taking the mean distance over 10 replicates). The Kendall-Colijn (KC) metric provides the greatest discrimination and is well defined for all tree topologies, and so we use this metric exclusively in subsequent analyses. At low mutation rates, there is a notable difference between the accuracy of `tsinfer`’s inferred trees when we use the standard KC metric and when we randomly break polytomies before measuring the distance between the trees. This is because there is little information available to resolve the nodes, and generating a random binary subtree on average results in something that is further from the truth than the original polytomy. Thus, `tsinfer`’s innate strategy of using polytomies to indicate uncertainty in a principled and systematic way has a significant advantage over methods that always fully resolve trees.

To evaluate the sensitivity of `tsinfer` to changes in the underlying simulation model, we also tested accuracy on more complex simulations. In Fig. S6 we show results of simulations of the three population Out-of-Africa model of human demography [Gutenkunst et al., 2009]. In this case, `tsinfer` does not seem to be affected by the underlying population structure and is a little more accurate than `ARGweaver` (although `ARGweaver` is less affected by error). In Fig. S7 we show inference accuracy on a simulated selective sweep, where we performed forward time simulations using simuPop [Peng and Kimmel, 2005] and ftprime [Kelleher et al., 2018]. In this case, `tsinfer` is substantially more accurate than the other tools after the advantageous mutation has swept to a reasonable frequency and also for many generations after fixation. In Fig. S8 we show the effect of running inference on a subset of the available haplotypes on `tsinfer`’s accuracy. We see that in the absence of error, having extra samples has little effect on the accuracy of inferences, but that larger samples can potentially help to correct for the presence of genotyping errors. This provides additional justification for using large sample sizes for ancestral inference.

To evaluate the computational performance of the different inference methods we measured the total user time and maximum memory usage (taking the mean over replicates). All experiments were run on a server with two Xeon E5-2680 CPUs and 256GiB of RAM. Fig. S9 shows the CPU time required for all four tools for varying sample sizes. The disparity in the running times is too large to show on a single scale, and `tsinfer` and `fastARG` are many times faster than `ARGweaver` and `Rent+`. Fig. S10 compares `tsinfer` and `fastARG` at a much larger scale, where `tsinfer` shows far better scaling in terms of CPU time and memory when increasing both sequence length and sample size. Empirically, `tsinfer`’s running time grows approximately linearly with sample size and super-linearly with sequence length on simulated data (Fig. S10).

All code for running the evaluations, including the precise version of each tool used, is included in the accompanying GitHub repository (https://github.com/mcveanlab/treeseq-inference/).

### Genealogical nearest neighbours

In our analysis of human data sets we use the genealogical nearest neighbours (GNN) statistic, which we define here. Let 𝕋 be a tree sequence where each tree *t* ∊ 𝕋 covers a length *L^t^* units of genetic sequence. Define *L* = Σ_*t*∊𝕋_ *L^t^*. Let *R* be a list of *K* sets of “reference nodes”, and let 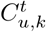 be the number of nodes ancestral to (and including) *u* in tree *t* from the set *R_k_*, with CU 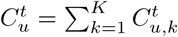. Then define

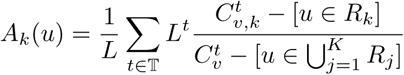

where *v* is the first node in *t* on the path to root from *u* such that 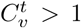, and [x] is an Iversonian condition such that [x] = 0 if x is false and [x] = 1 otherwise.

### Empirical data preparation

All human data used is relative to the GRCh37 reference. We used ancestral states from Ensembl release 75, as this is the most recent version available for GRCh37. For each data set we processed the downloaded variation data to produce a .samples file used as input to `tsinfer`. In this preprocessing step we kept only biallelic SNPs with a high-confidence ancestral state and with phased calls for all samples. Any singleton or invariant sites were discarded as these provide no information about topology. Sites with frequency (*n* — 1) were also discarded as these are likely to be highly enriched for miscalled ancestral states. Details of the data sources used and a fully automated pipeline for obtaining public data sets, preprocessing the variation data and constructing the tree sequences are included in the accompanying GitHub repository (https://github.com/mcveanlab/treeseq-inference/).

### Outlier samples

We observed two particularly large outliers when considering the number of sample edges per individual in the TGP data set: NA20289 from the ASW population and HG02789 from PJL (see Figs. S13 and S14). Both parental nodes for NA20289 have roughly double the number of sample edges as the population mean. The physical position of the breakpoints of the parental nodes of NA20289 seemingly reflects an abundance of phasing switch errors, potentially explaining this finding. At the location of a phasing switch error, the parental nodes of a sample would swap the ancestors they were copying from, resulting in a far greater number of sample edges than if phasing error had not occurred. When considering all TGP samples, the average paternal and maternal node breakpoints were within 100 base pairs of each other only 32 times in the unsimplified tree sequence, while in NA20289 this occurred approximately 551 times. HG02789 showed only 30 breakpoints within 100 base pairs of one another, which is unsurprising since only one parental node exhibited a high number of sample edges.

In the SGDP data set we observed the S-Naxi-2 had a highly elevated sample edge count. Since this sample also had no associated metadata, we removed it from the analysis.

### Duplicate UKB/TGP samples

We observed 9 individuals in the UKB data set that are likely to also be in TGP. Using the (unsimplified) UKB+TGP tree sequence, we first found outliers with low numbers of sample edges. Then, from the pool of samples with fewer than 50 edges where they are children, we extracted those for which both nodes in an individual were present (i.e., both maternal and paternal nodes have unusually low sample edge counts). This left a total of 9 candidate individuals; for 8 of these, both nodes of the UKB individual copied from a single node of a TGP individual over 97% of the sequence length (mostly > 99%). The nodes in the UKB and TGP individuals paired up exactly, signifying that the phasing for these individuals is in agreement in the two data sets. In the 9th individual we observed very high copying (> 97%) from a single TGP individual, but with switches between the two nodes. Let *p*_1_ and *p*_2_ are the parental nodes in the TGP individual; we observed 61% copying from pi and 37% from *p*_2_ in the first node of the UKB individual, and 60% copying from *p*_2_ and 38% from *p*_1_ in the second. This is likely to indicate phasing errors for this individual in one of the data sets.

This analysis required 42 seconds of CPU time to run using simple Python code.

## Acknowledgements

**Acknowledgements**

This work was supported by the Wellcome Trust [100956/Z/13/Z]. We would like to thank John Novembre and Peter Ralph for helpful comments on earlier drafts of this manuscript. Thanks also to Peter Ralph and Kevin Thornton for many useful discussions on tree sequence algorithms.

## Availability

All code used to run evaluations in this paper is available at https://github.com/mcveanlab/treeseq-inference. `Tsinfer` is freely available under the terms of the GNU GPL; see the documentation at https://tsinfer.readthedocs.io/ for further details.

## Supplementary material

**Figure S1:**
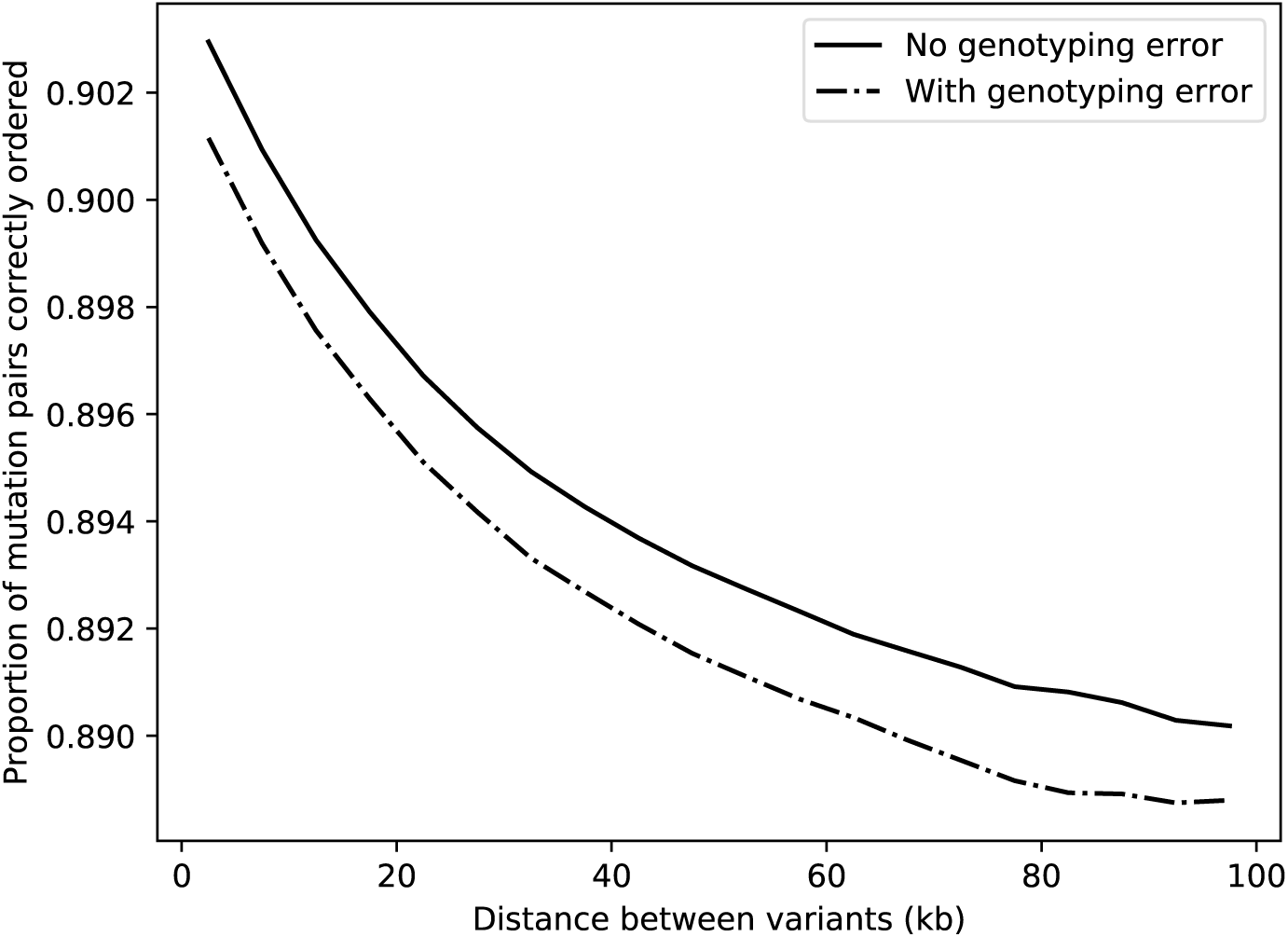
Accuracy of frequency as a proxy for relative age. 50 sample sequences of 100kb were simulated using `msprime` with *N*_e_ = 5000 and *μ* = *ρ* = 10^−8^. For all pairs of variant sites, the difference in mutational age was compared to the difference in frequency between the two derived alleles. If *T*(*a*_1_) and *T*(*a*_2_) are the ages of the derived alleles at sites 1 and 2 respectively, and *f* (*a*_1_) and *f* (*a*_2_) are their current frequencies in the population, frequency is said to correctly predict age order if *T*(*a*_1_) < *T*(*a*_2_) == *f* (*a*_2_) < *f* (*a*_2_). We show the proportion of pairs for which frequency provides the correct order of mutation age. Results were binned into 5kb intervals and averaged over 250,000 replicates. Correlation of adjacent tree topologies results in frequency being a better predictor of relative age when variants are physically close to one another. Note that singletons have been included when calculating the statistics in this figure; removing singletons (which are not used for `tsinfer` inference) results in an average accuracy roughly 5% lower than shown here.

**Figure S2:**
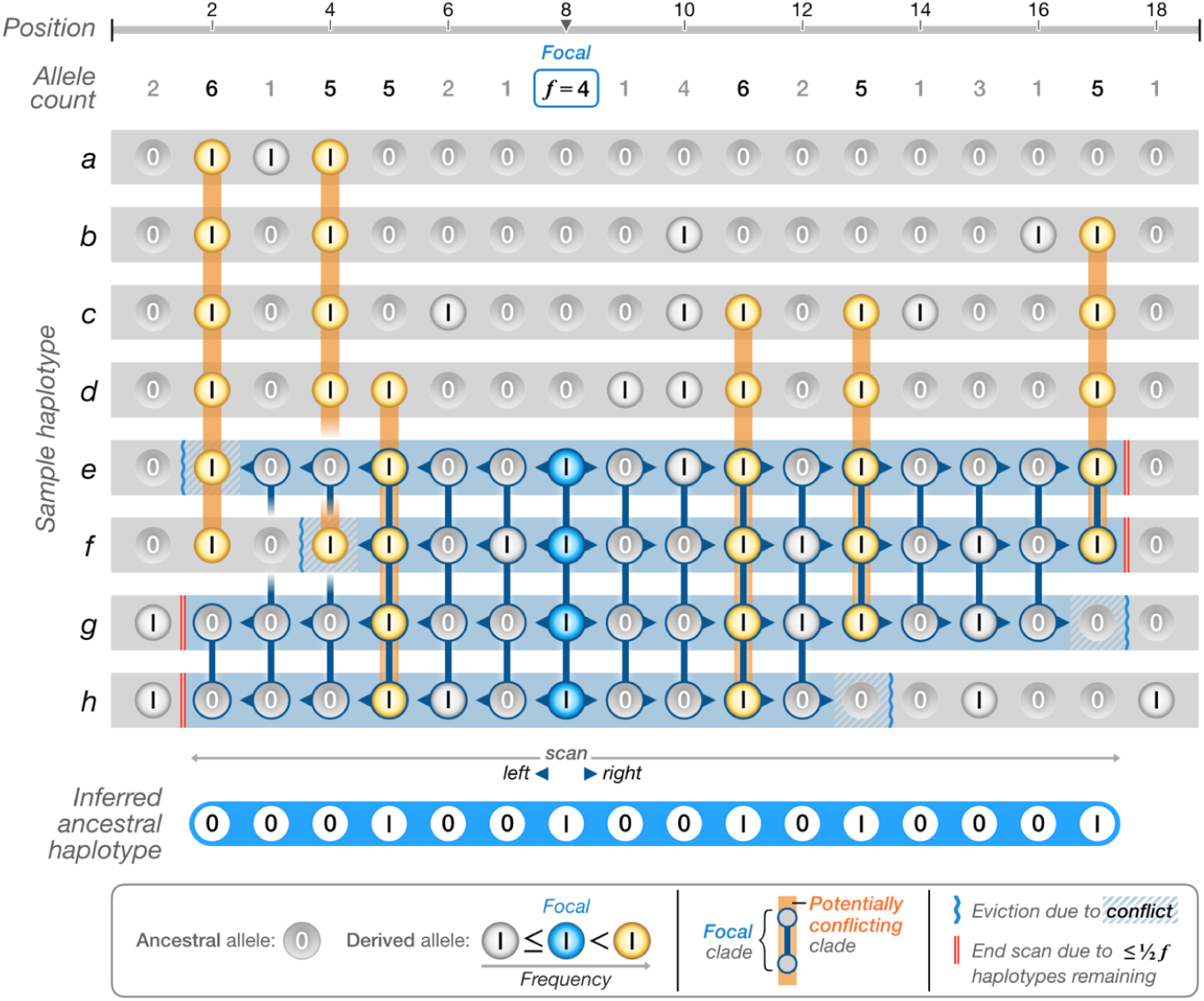
Schematic of ancestral haplotype reconstruction. We are constructing an ancestral haplotype for the ancestor corresponding to site 8. The time of each site is approximated by its corresponding frequency value (referred to as allele count here). Note that this is a simplified schematic: the actual implementation includes a dampening effect such that haplotypes are evicted from S if they disagree with the consensus call at two adjacent sites.

**Figure S3:**
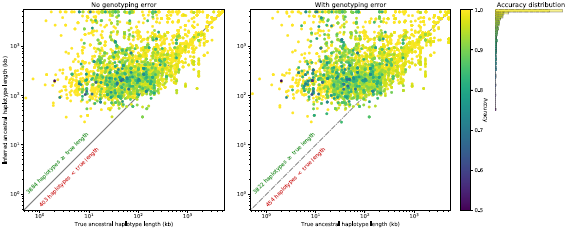
Accuracy of inferred ancestral haplotypes. True ancestral haplotypes were obtained from a single simulation of 100 sample sequences, each of 5Mb in length, using `msprime` with *N*_e_ = 5000, and *μ* = *ρ* = 10^−8^. A typical result is shown here, with all ~ 4300 separate ancestral haplotypes plotted as individual points. Accuracy measures the fraction of correctly reconstructed sites, the calculation being restricted to the region in which the true and reconstructed haplotypes overlap.

**Figure S4:**
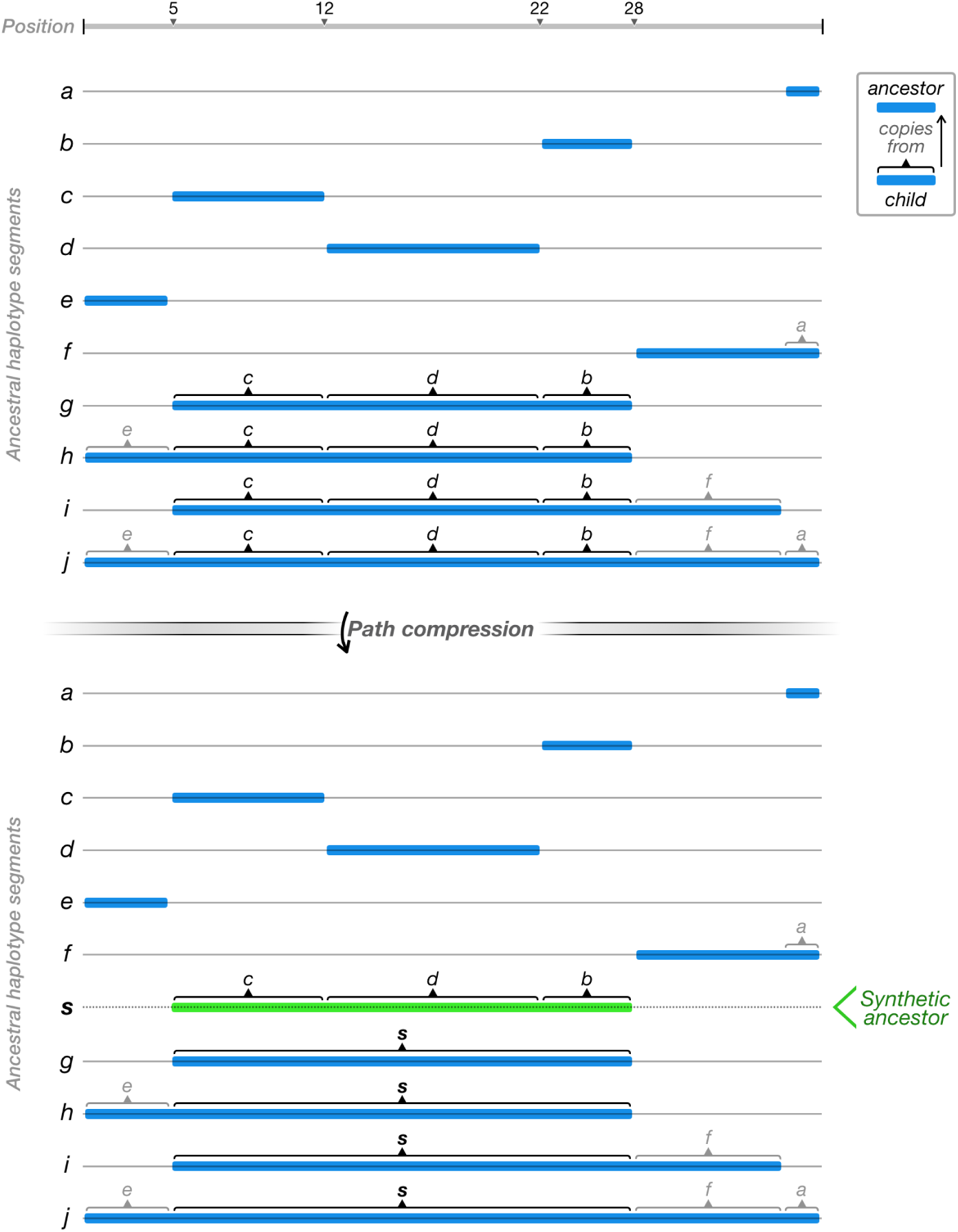
Path compression schematic. In the top panel, the children *g*, *h*, *i* and *j* copy from ancestors above them (*a* through *f*). Several sets of edges point to the same parent in the same genomic location, but from different children (e.g. the edges from *g* → *c*, *h* → *c*, *i* → *c*, and *j* → *c*). If there are multiple such sets adjacent to each other, the number of edges can be reduced by inserting a synthetic ancestor *s*, to which the children all point (bottom panel). This compression of ancestral paths essentially treats shared recombination breakpoints as evidence of shared ancestry.

**Figure S5:**
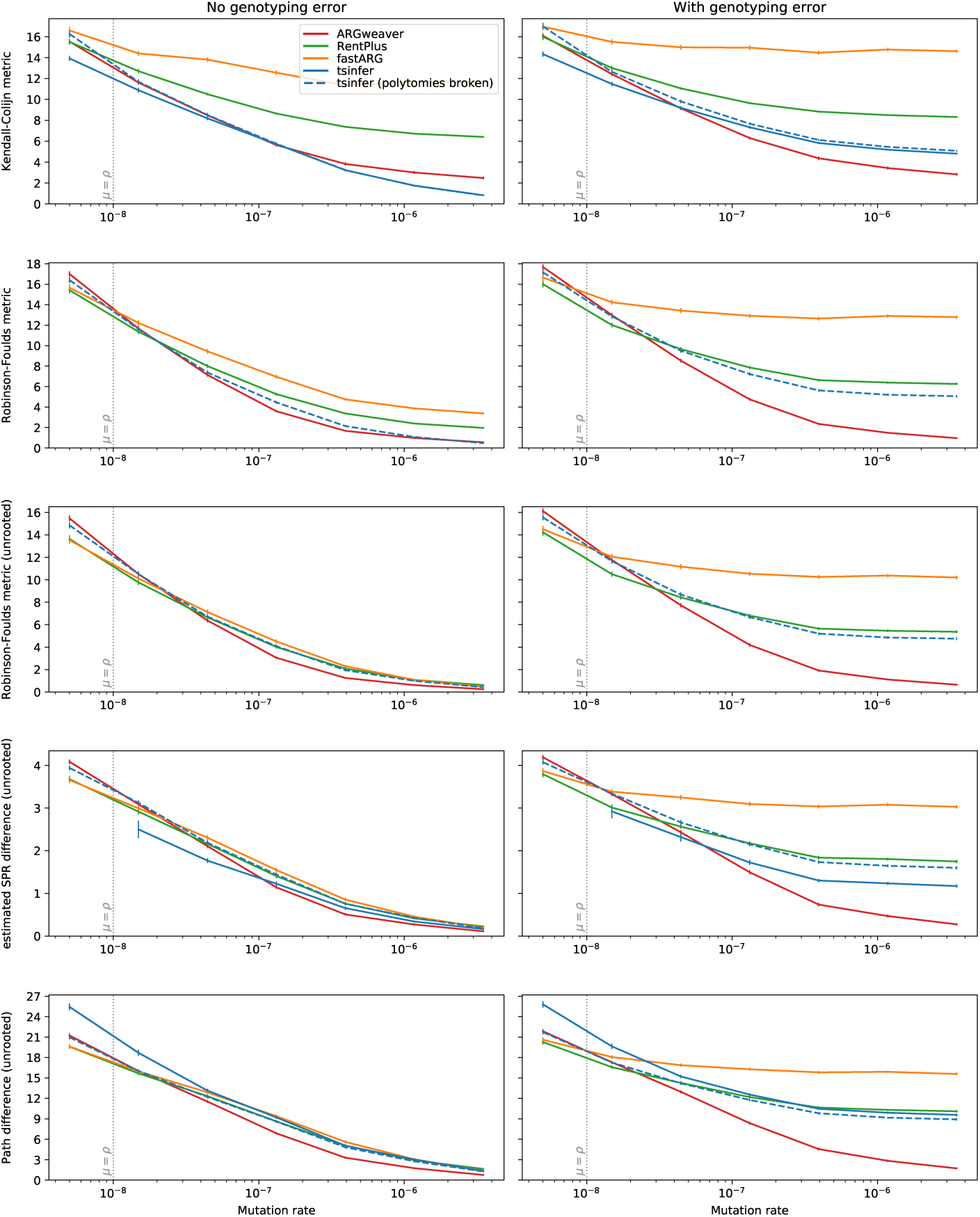
Accuracy of ancestral inference according to different tree distance metrics. 16 sequences were simulated, each of 100kb in length, using `msprime` with *N*_e_ = 5000 and *ρ* = 10^−8^. Simulations were replicated 100 times for each of seven different mutation rates. The region corresponding to human-like parameters, where *μ* ≈ *ρ*, is marked by a vertical dotted line.

**Figure S6:**
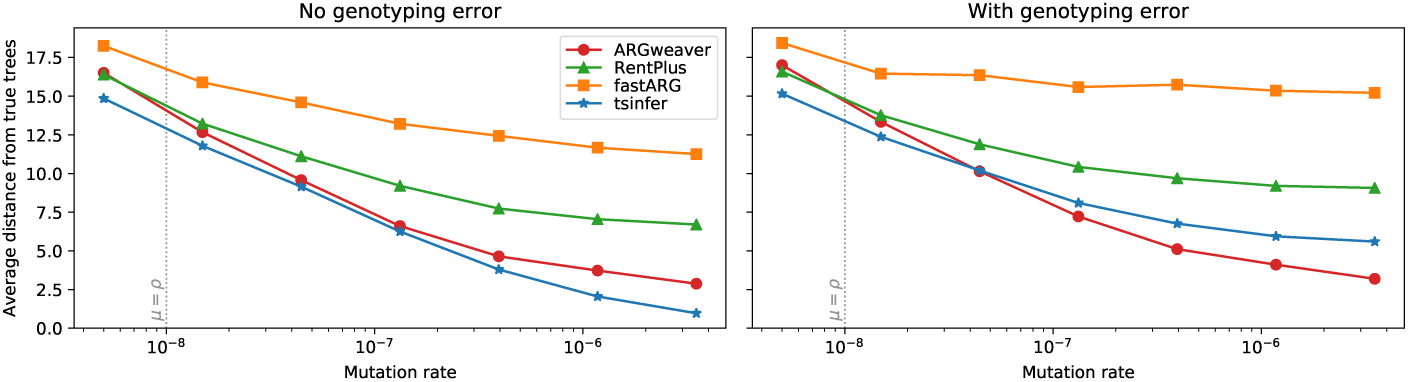
Accuracy of inference for different tools under a three-population Out-of-Africa model, as implemented in `msprime`. 16 sample haplotypes (6 African, 6 European, and 4 Asian) were simulated, each of length 100kb, with *ρ* = 10^−8^ under a variety of mutation rates. Each point represents an average over 100 replicates.

**Figure S7:**
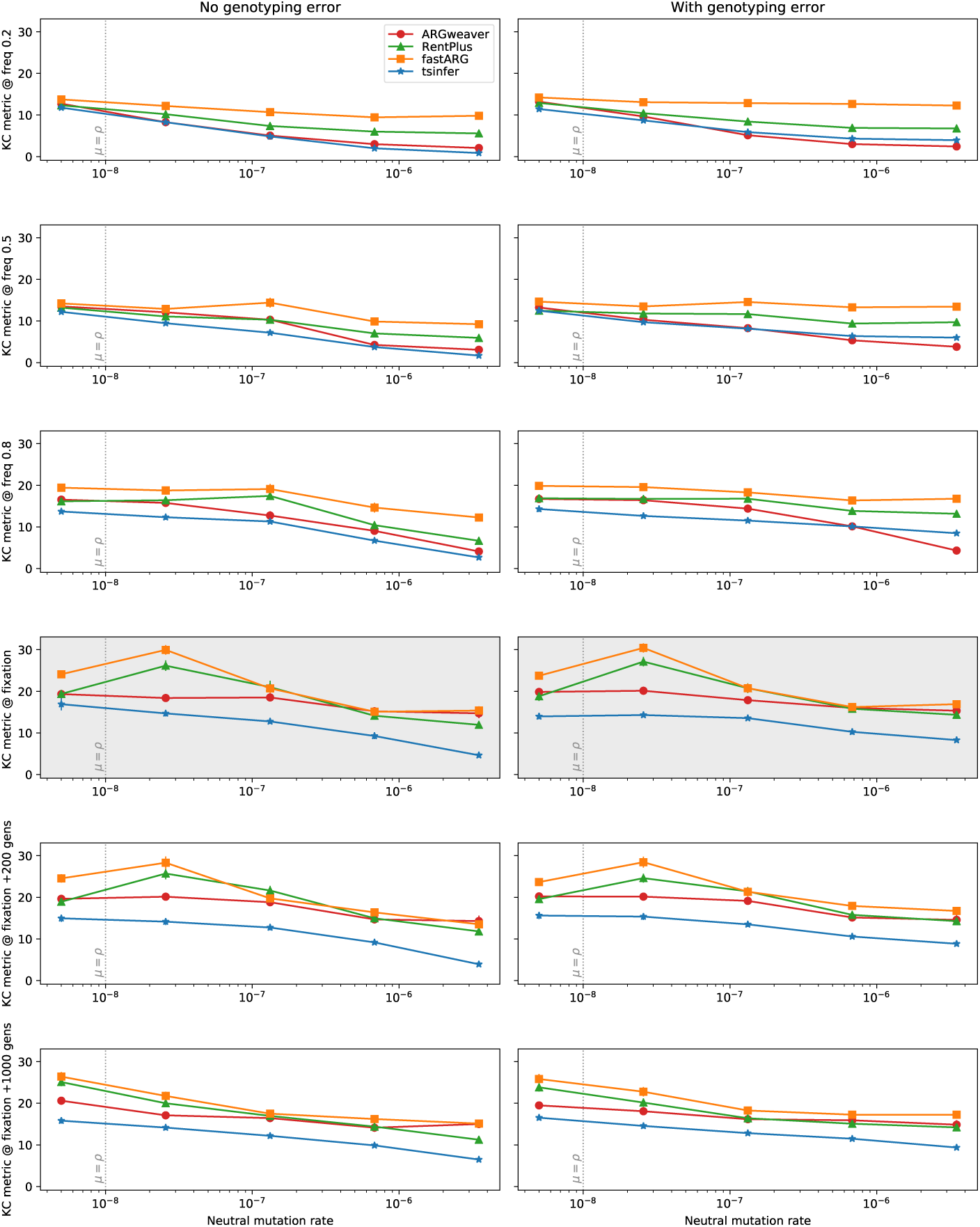
Accuracy of inference at different points during and after a selective sweep. Forward-time simulations were performed for sequences of length 100kb, in a Wright-Fisher panmictic population of size *N*_e_ = 5000, with a neutral mutation rate of *μ* = *ρ* = 10^−8^. A single advantageous mutation in the middle of the sequence was introduced and tracked until it went to fixation (or reintroduced if it became extinct). This allele was associated with a selection coefficient of *s* = 0.1 in homozygotes, with a dominance coefficient of *h* = 0.5. Inference was performed for 16 sample sequences when this allele first hit a frequency of 20, 50, and 80%. The situation at fixation is highlighted in grey: lower rows show inference accuracy 200 and 1000 generations post-fixation. Each point is an average over 50 simulations.

**Figure S8:**
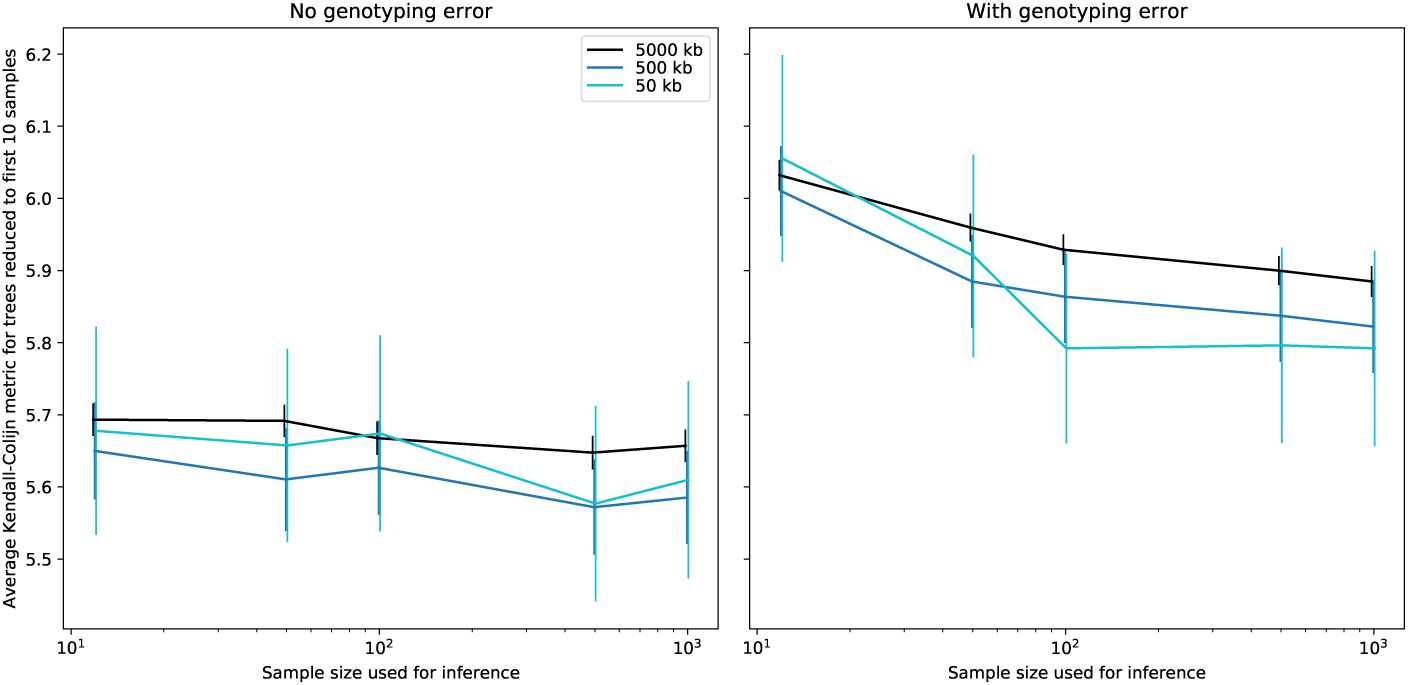
Effect of increased sample size on inference accuracy. Neutral simulations were replicated 100 times for each of three different sequence lengths using `msprime` with *N_e_* = 5000 and *μ* = *ρ* = 10^−8^, and a large initial sample size of 1000. To allow like-for-like comparison, the tree distance metric was restricted to comparisons involving the simulated and inferred ancestral topologies of the first 10 samples only. Tree sequence inference was based on the first 12, 50, 100, 500, or 1000 samples, out of the total simulated set of 1000 haplotypes.

**Figure S9:**
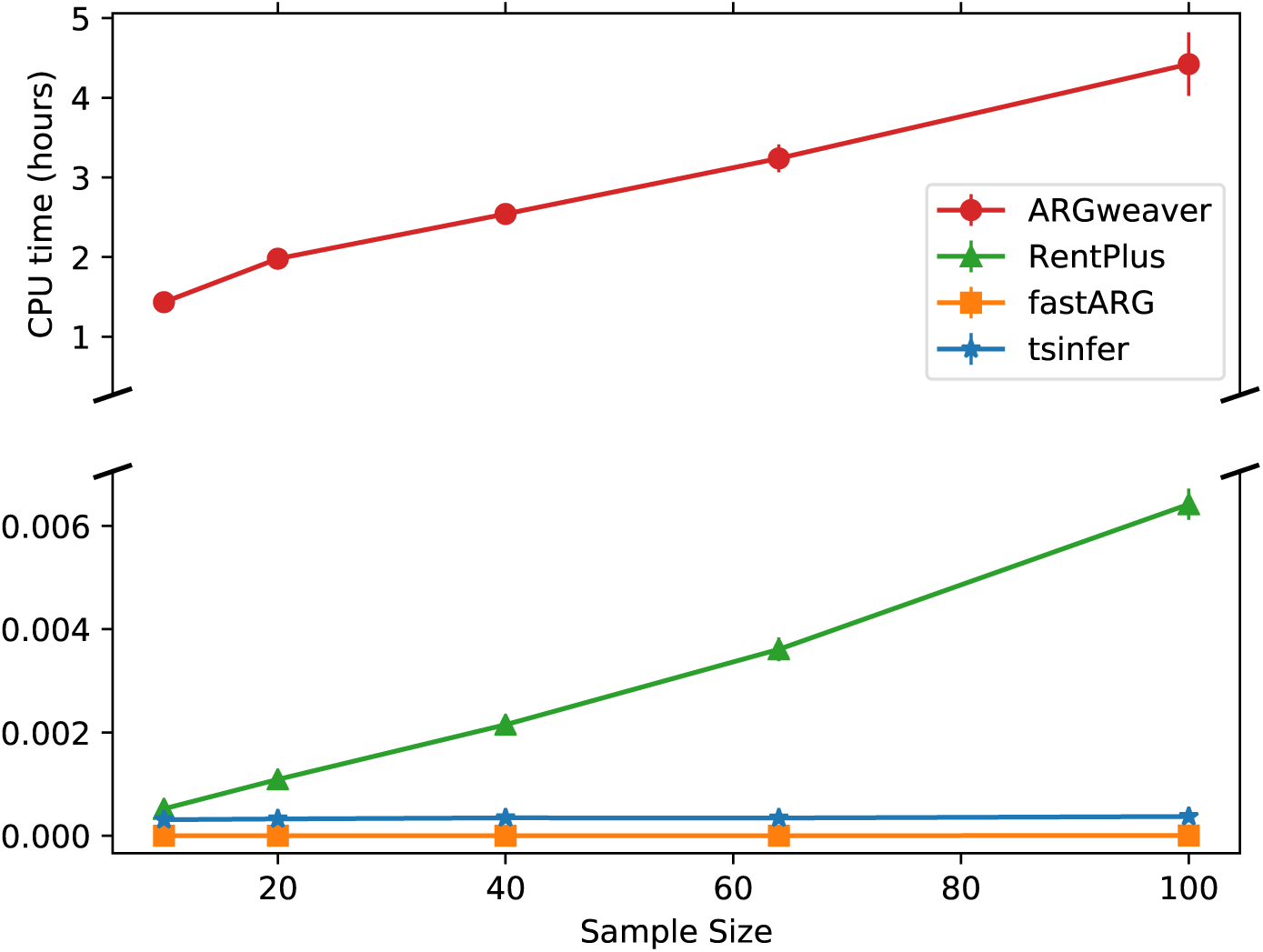
CPU time for ancestral inference as a function of sample size.Sequences of 200kb in length were simulated using `msprime` setting *N_e_* = 5000 and *μ* = *ρ* =1 × 10^−8^, then used for inference without imposing genotyping error. Each point represents an average over 10 replicates. `ARGweaver` requires over 3 orders of magnitude more CPU time than the other methods, hence a discontinuous y-axis is used to fit all tools onto the plot.

**Figure S10:**
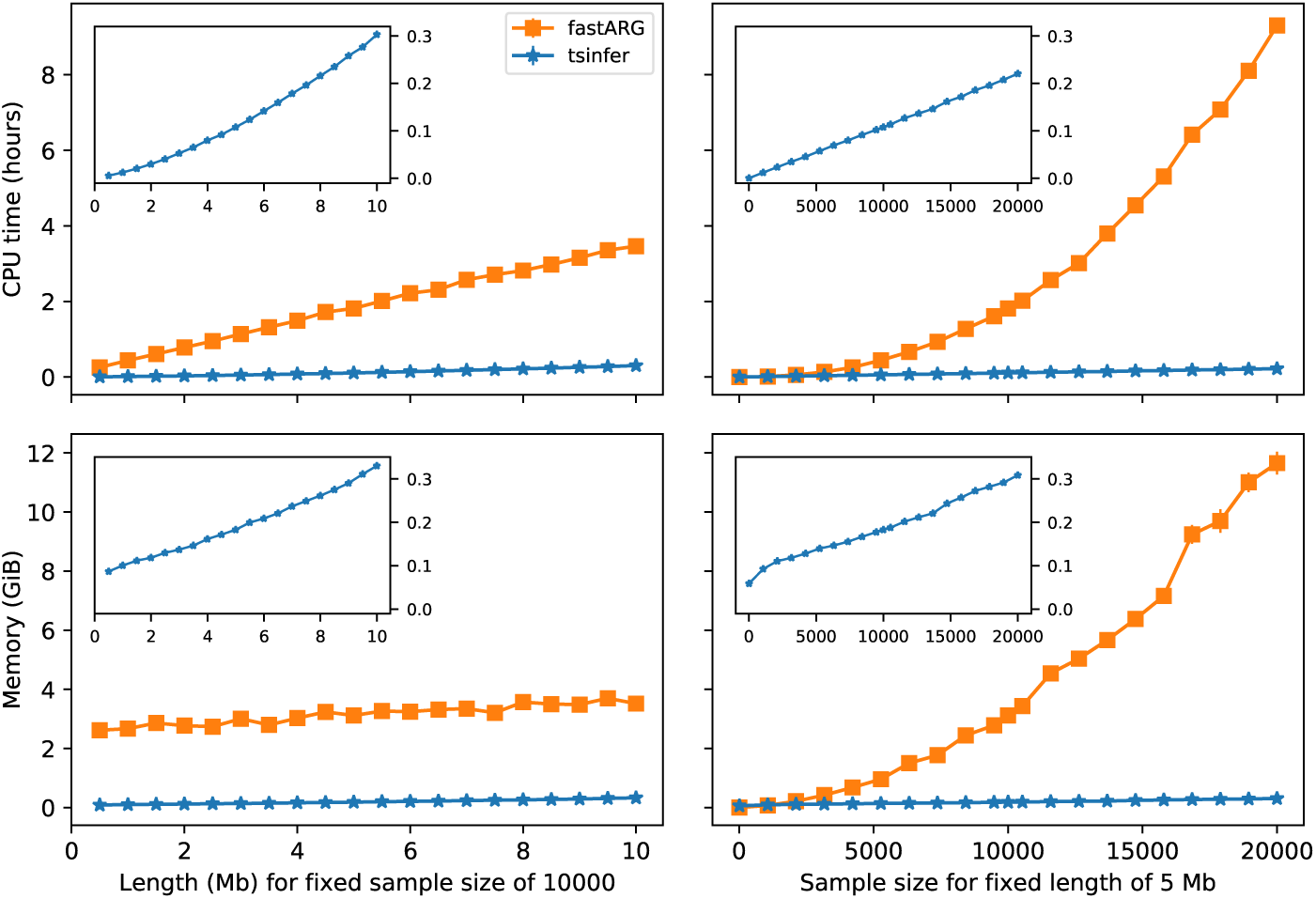
CPU time and memory requirements for `tsinfer` and `fastARG`, as a function of sample size and sequence length. Sequences were simulated using `msprime`, setting *N_e_* = 5000 and *μ* = ρ =1 × 10^−8^, then used for inference without imposing genotyping error. Each point represents an average over 50 replicates. The scaling properties of `tsinfer` are hard to examine using the same y-axis scale as for `fastARG`, hence the inset plots show the `tsinfer` line only, with a rescaled y-axis.

**Figure S11:**
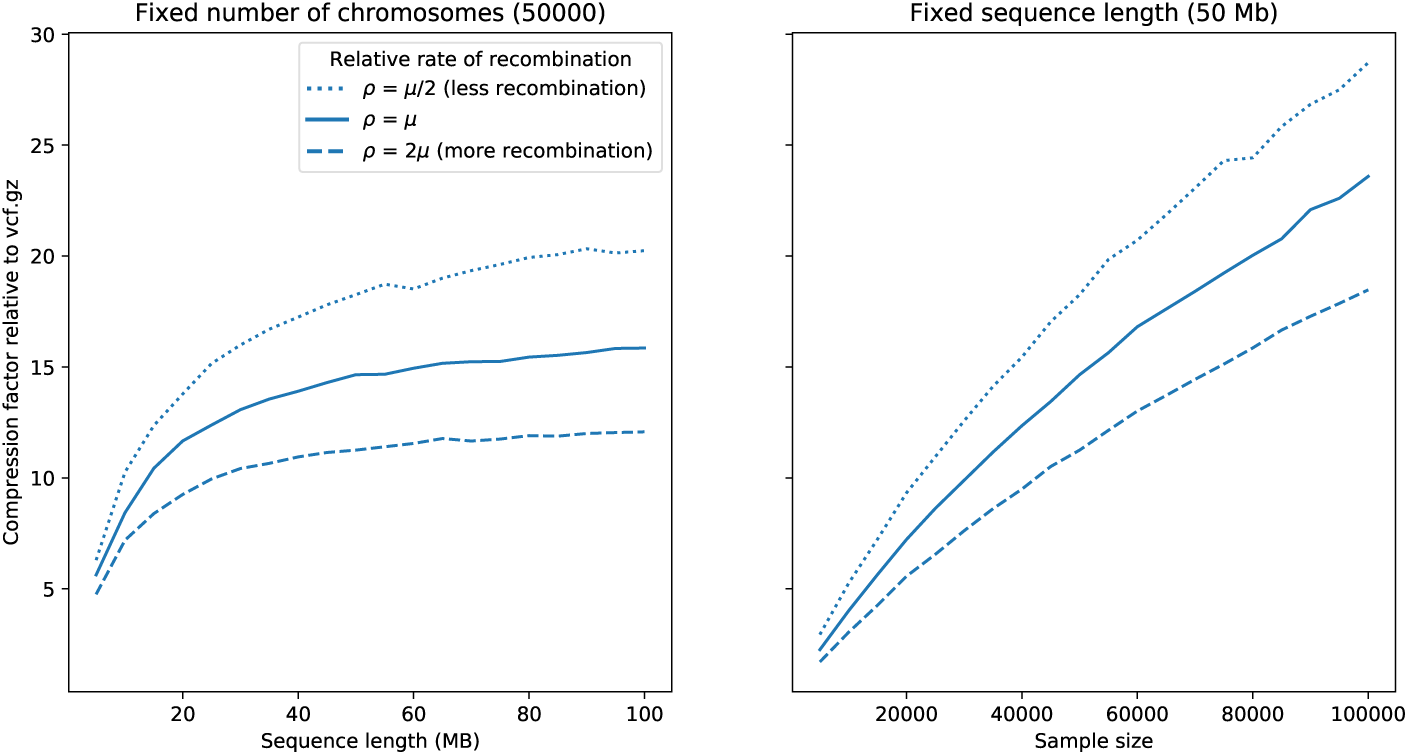
Mean compression factor achieved by `tsinfer` compared to gzipped VCF data for varying sequence lengths and sample sizes, under three different recombination rates. Sequences were simulated using `msprime`, setting *N_e_* = 5000 and *μ* = 10^−8^, then used for inference without imposing genotyping error. Data are averages over 10 replicates for each combination of parameters.

**Figure S12:**
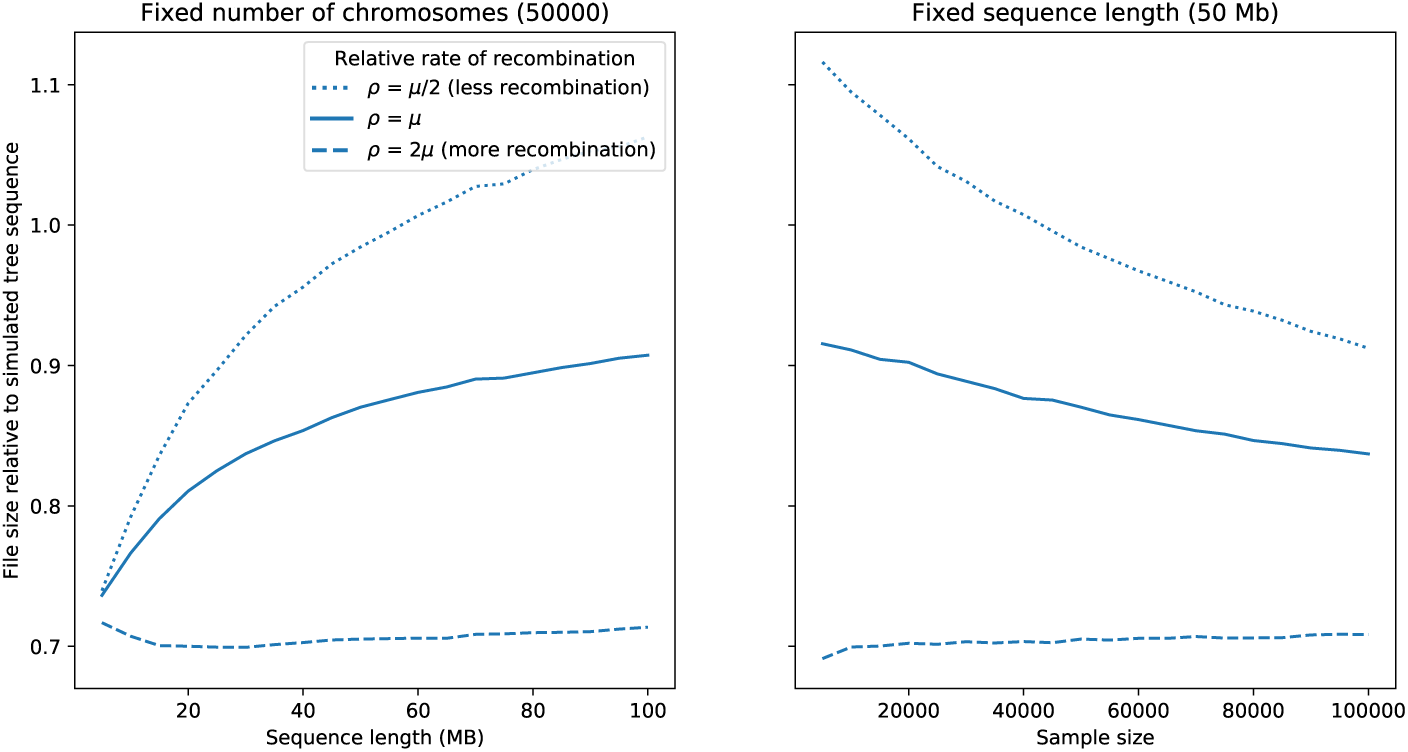
The relative filesize of tree sequences inferred by `tsinfer` relative to the original tree simulated sequence, for varying sequence lengths and sample sizes, under three different recombination rates. Sequences were simulated using `msprime`, setting *N_e_* = 5000 and *μ* = 10^−8^, then used for inference without imposing genotyping error. Data are averages over 10 replicates for each combination of parameters.

**Figure S13:**
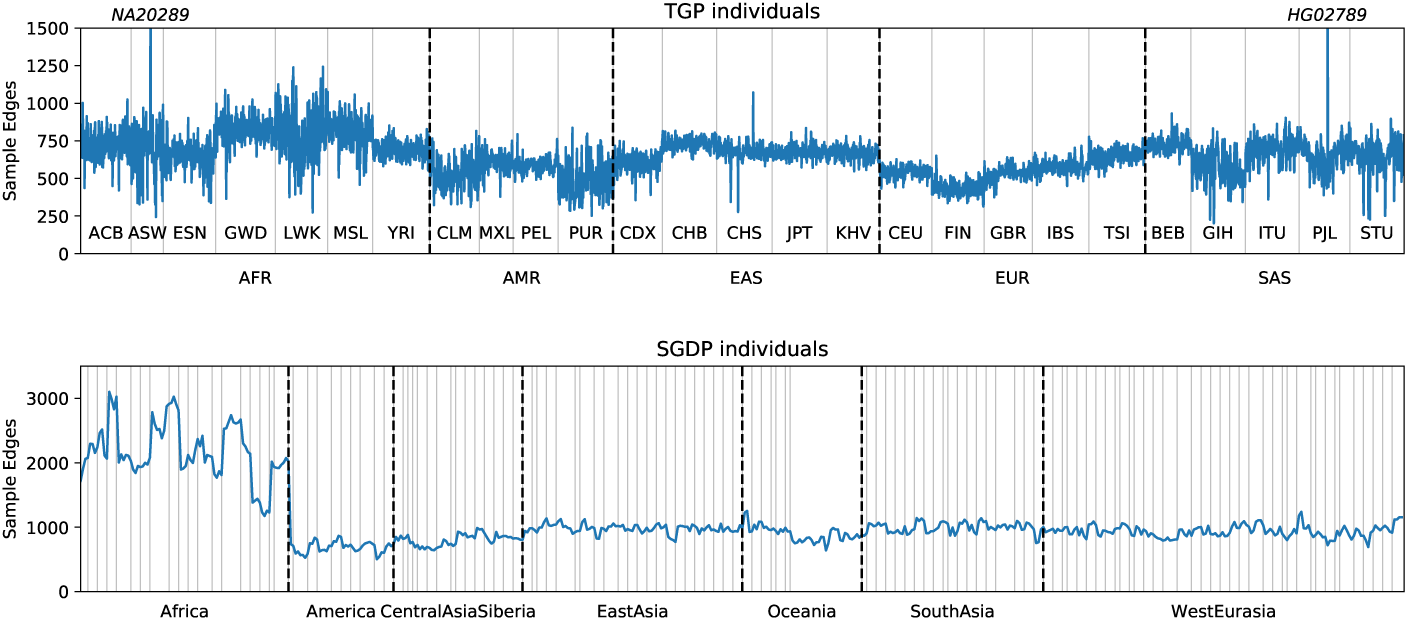
Summary of sample edges across all TGP and SGDP populations. The number of edges per sample for all individuals in TGP and SGDP, organised by population and continent.

**Figure S14:**
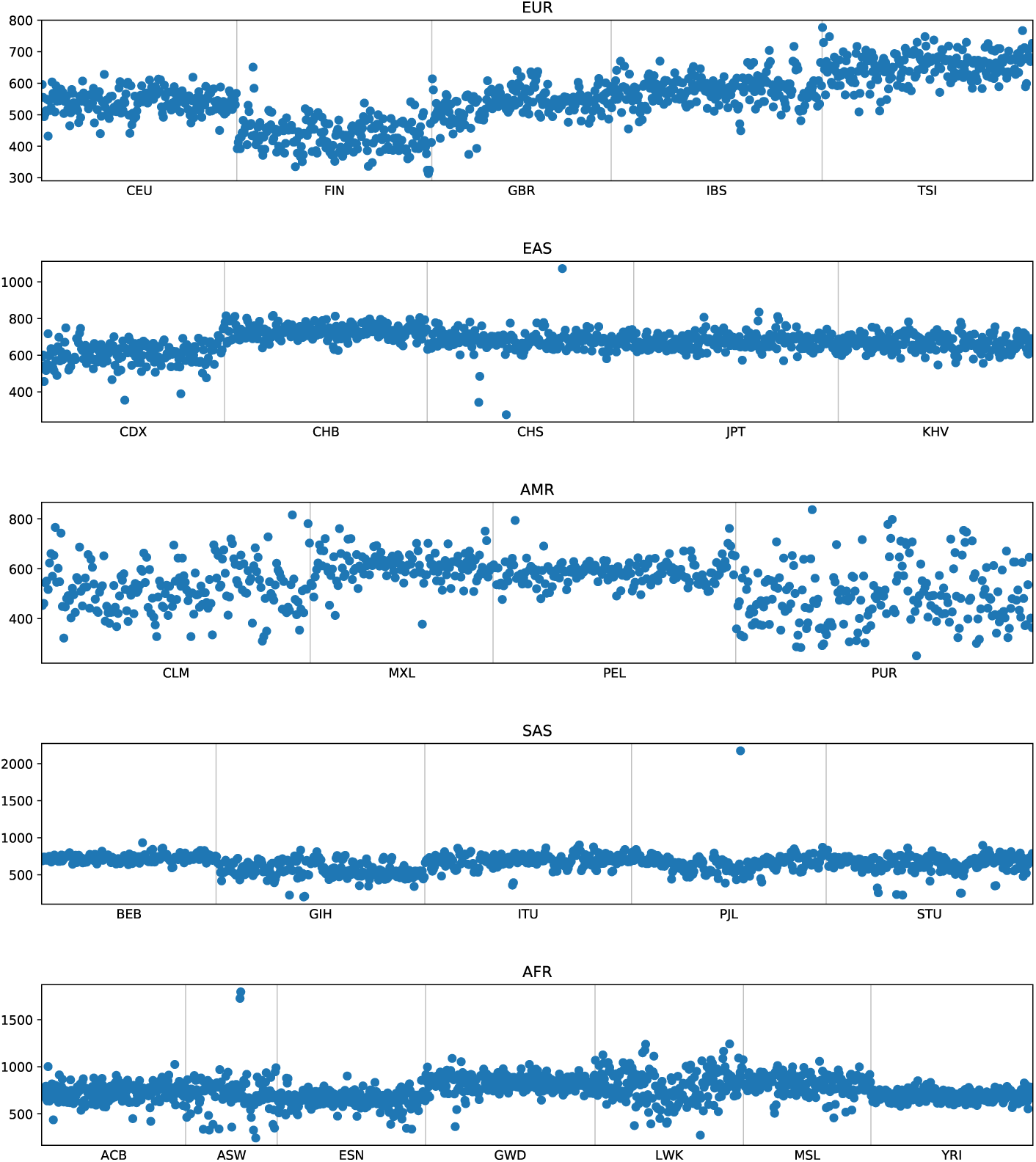
Sample edges for the TGP populations. The number of edges per sample for all individuals in TGP organised by population, with each continent plotted separately.

**Figure S15:**
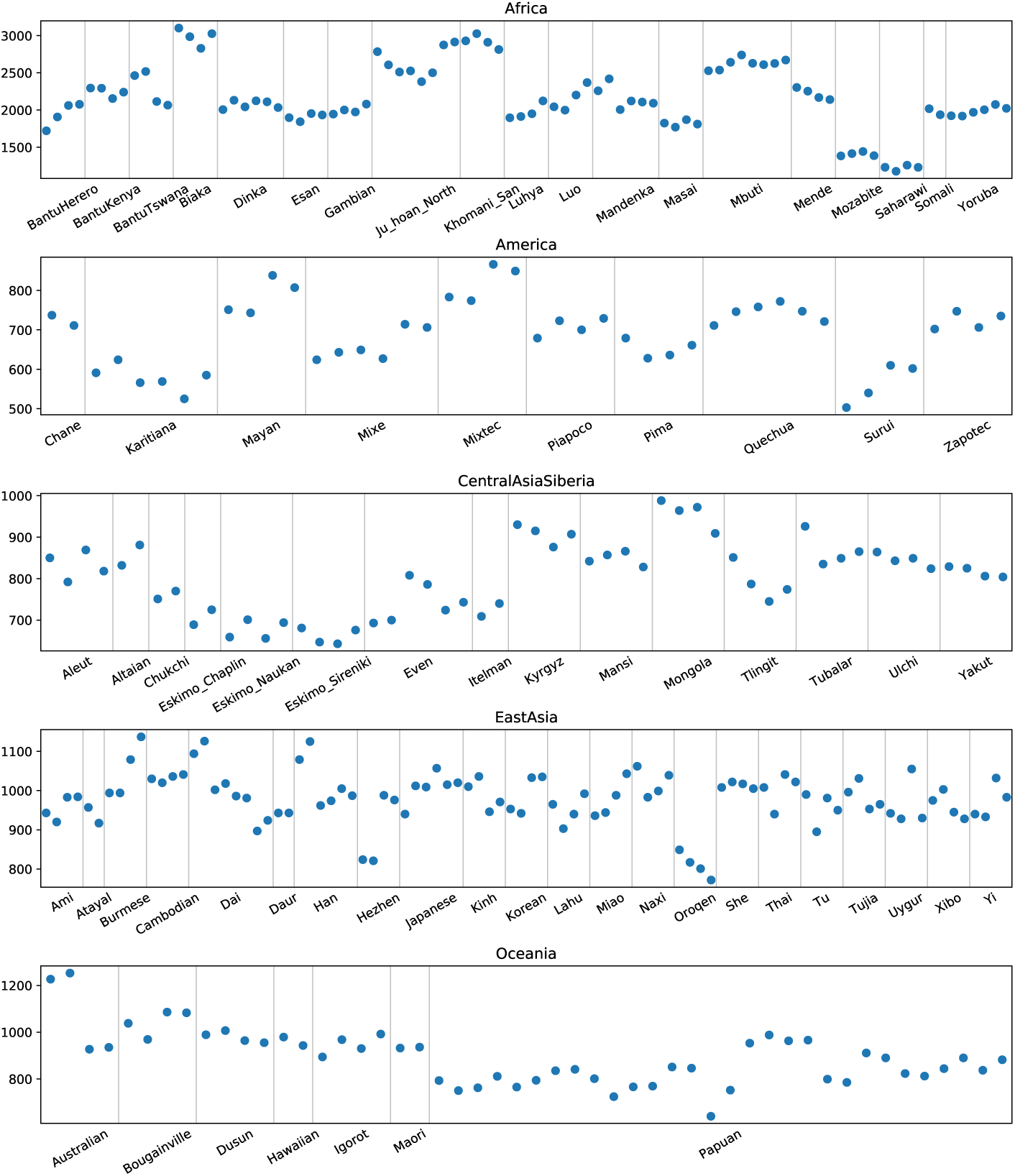
Sample edges for the SGDP populations. The number of edges per sample for all individuals in SGDP organised by population, with each continent plotted separately.

**Figure S16:**
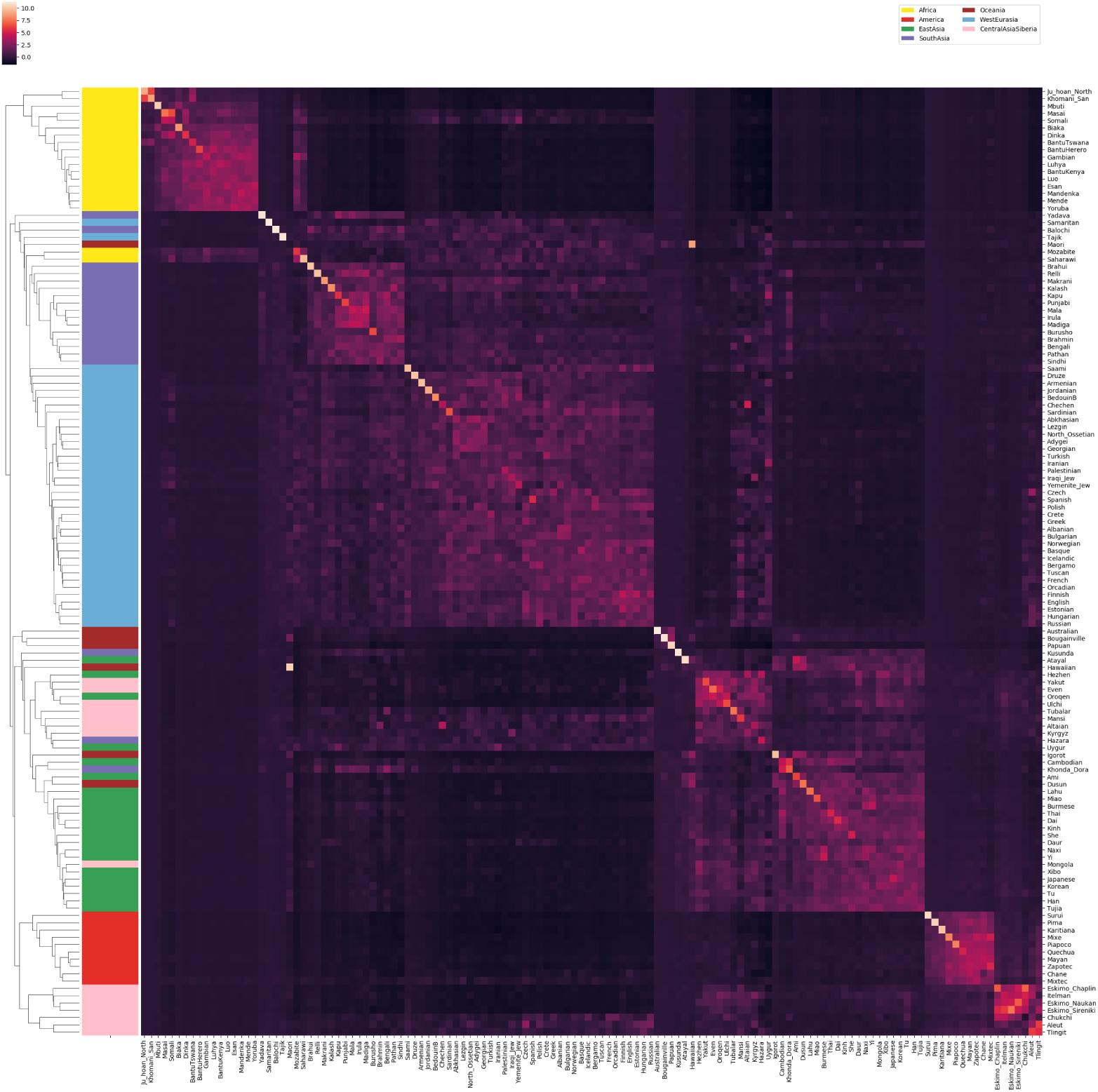
Z-score normalised GNN proportions for SGDP by population. The GNN matrix was first z-score normalised by column and the rows then hierarchically clustered.

